# Long-term Epstein-Barr virus persistence in immunocompetent, primate-related Chinese tree shrews and inducible lytic reactivation in species-matched epithelial cultures

**DOI:** 10.64898/2026.11.11.765928

**Authors:** Janardhan Avilala, Hirotomo Dochi, Joseph Wekselblatt, Erik K Flemington, Amy Pierce, Christopher Roger Brown, Georgina Dobek, Kylie Ott, Paige Roussel, Ramsy Abdelghani, David Becnel, Mark Sides, Gilbert Morris, Hong Liu, Wenke Feng, Zhen Lin

**Author notes:** These authors contributed equally (Janardhan Avilala, Hirotomo Dochi).

## Abstract

Experimental models combining long-term Epstein-Barr virus (EBV) infection in immunocompetent animals with species-matched cellular systems for studying viral regulation remain limited. Chinese tree shrews combine evolutionary proximity to primates with small-animal practicality and native immunocompetence, offering a complementary nonrodent model. We serially monitored blood and oral specimens from six animals inoculated with Akata EBV. Five were followed for approximately two years, including two beyond two years, and the sixth underwent tissue collection at approximately six months. Intermittent EBV DNA detection included blood positivity beyond two years in one animal. Viral RNA localized to liver, lung, and spleen at six months and to liver and spleen of another animal near two years, supporting persistent infection. All tested blood and oral specimens from six uninoculated controls were negative. In parallel, renewable lung-derived epithelial/mesothelial-like cultures maintained EBV infection in two immortalization backgrounds. Viral profiling identified Qp-associated EBNA1 transcription and latency-associated expression alongside spontaneous lytic transcription. Host transcriptomics identified 684 robust EBV-associated expression changes, with prominent enrichment of MYC and E2F target programs and metabolic pathways, together with reduced interferon- and inflammatory-response transcription across both backgrounds. These findings nominate MYC/E2F-associated host programs as candidate regulators of the balance between maintained infection and lytic activity. Zta expression and sodium butyrate induced lytic reactivation, with infectious virus production demonstrated after sodium butyrate treatment. This integrated platform complements humanized mice and nonhuman-primate models and connects long-term EBV persistence in vivo with experimentally accessible studies of host regulation and the latent-to-lytic transition in species-matched cultures.

**IMPORTANCE:** Epstein-Barr virus (EBV) persists for life and contributes to cancer, but models connecting prolonged infection with viral regulation remain limited. Chinese tree shrews combine evolutionary proximity to primates with small-animal practicality and a native immune system without human-cell engraftment. Serial monitoring of six inoculated animals identified blood viral DNA beyond two years in one animal and late tissue viral RNA in another, supporting persistent infection. Renewable cultures from the same species maintained latency-associated expression and supported induced reactivation and infectious virus production. Importantly, infected cultures showed enrichment of MYC and E2F target programs alongside reduced immune-related transcription. These findings identify a host-expression state that provides a mechanistic framework for investigating how cellular growth programs influence the balance between EBV latency and reactivation. By integrating prolonged animal infection with species-matched cultures, this platform complements humanized mice and nonhuman-primate models and enables investigation of host pathways that may sustain persistent infection.

## INTRODUCTION

Epstein-Barr virus (EBV) is a ubiquitous gammaherpesvirus that establishes lifelong infection in more than 90% of adults and contributes to infectious mononucleosis, lymphoid and epithelial malignancies, and immune-mediated disease such as multiple sclerosis (1-5). Humans are the natural host for EBV, and conventional mice do not reproduce authentic EBV infection. Hematopoietic humanized mice have provided important insights into primary infection, immune control, and EBV-associated lymphoproliferation, but they remain chimeric and variably reconstituted and do not routinely provide a human epithelial compartment alongside the human hematopoietic compartment (6, 7). Rhesus macaque studies commonly use host-adapted lymphocryptoviruses rather than EBV itself. Although these models capture infection in a natural primate host, antigenic and biological differences can complicate direct evaluation of EBV-specific interventions (8).

The Chinese tree shrew (*Tupaia belangeri chinensis*) is a small, immunocompetent nonrodent in the order Scandentia. Tree shrews are evolutionarily closer to primates than rodents while retaining practical features of small laboratory animals, including compact body size, comparatively short reproductive cycles, and lower maintenance requirements than larger nonhuman primates (9, 10). These features make tree shrews attractive for studies requiring repeated sampling and prolonged follow-up. For EBV research, infection in a native host environment without human hematopoietic engraftment offers a distinct complement to humanized mice and primate lymphocryptovirus systems.

Wang et al. demonstrated intermittent EBV DNA detection and viral markers in lymphoid tissues after experimental challenge, with observation periods extending to 21 weeks (11). Xia et al. documented blood and oral viral DNA, tissue infection, CR2-associated receptor features, and proliferation of EBV-exposed peripheral blood mononuclear cells (PBMCs), following infection through 91 days (12). Subsequent studies examined viral reactivation (13) and the comparative infectivity of B95-8 and P3HR-1 laboratory strains over a 28-day period (14). These studies established susceptibility, but serial monitoring over a substantially longer interval is needed to evaluate whether experimentally introduced EBV remains detectable beyond the initial months of infection.

An important consideration in interpreting earlier tree shrew studies is viral strain. B95-8 EBV contains an approximately 12-kb deletion that removes a substantial portion of the BamHI A rightward transcript (BART) region, including multiple viral microRNAs (15, 16). P3HR-1 carries an EBNA2-associated deletion and is defective for conventional B-cell transformation (17). These laboratory strains are useful experimental tools, but strain-specific deletions complicate direct comparisons of host range, latency programming, and pathogenesis. To avoid these particular genomic deletions, we employed Akata GFP-EBV, a widely used GFP-expressing recombinant derived from the well-characterized Type I Akata strain (18). Unlike the deletion-bearing B95-8 and P3HR-1 laboratory strains, Akata GFP-EBV retains the BART region and intact EBNA2 locus, providing a more complete EBV genomic background for investigating viral infection, persistence, and latency.

The epithelial compartment is important in EBV biology. EBV persists and replicates at oral and nasopharyngeal mucosal sites and is linked to nasopharyngeal and gastric carcinomas. EBV has also been detected in a subset of lung cancers (3, 4). EphA2 is an epithelial entry receptor in human cells (19, 20), whereas CR2/CD21 provides a relevant receptor context for B-cell infection. A species-matched cellular system should address not only susceptibility but also the viral functions retained after infection is established. Combining such cultures with longitudinal animal observations would permit complementary assessment of persistence, promoter-associated transcription, host-expression states, lytic inducibility, and infectious output.

Here, we established a Chinese tree shrew colony and combined longitudinal blood and oral monitoring after Akata GFP-EBV inoculation with tissue EBER RNAscope at earlier and late collection points. Follow-up extended beyond two years in two animals, with blood EBV DNA detected beyond two years in TS05. The tissue analyses included TS06 at approximately six months and TS01 near two years. Uninoculated animals provided blood, oral, and tissue controls. In parallel, we established hTERT-and T-antigen-immortalized lung-derived cultures to examine viral transcription and the functions retained during maintained infection. Receptor characterization and lineage profiling defined the cellular context. Promoter-associated transcripts, host-expression programs, downstream viral proteins, and Raji-recipient assays then connected molecular characterization with lytic inducibility and infectious output. Together, these approaches combine a small, primate-related host with species-matched cultures for complementary studies of prolonged EBV persistence in vivo and viral regulation in vitro.

## MATERIALS AND METHODS

### Animals, husbandry, and ethics

Thirteen individual Chinese tree shrews (Tupaia belangeri chinensis, nine males and four females) were included in the study (**Table S2**). Six animals (three males and three females) were inoculated with Akata GFP-EBV at 11.6-38.9 months of age (mean ± SD, 24.6 ± 12.3 months). Five animals (TS01-TS05) formed the longitudinal cohort. TS01 also contributed tissues for late EBER analysis. TS06 was separately designated for serial monitoring and subsequent tissue analysis. Six uninoculated animals (NC01-NC06, five males and one female) provided blood and oral controls and were 17.5-27.8 months old at first sampling (19.4 ± 4.1 months). An additional uninoculated male, NC07, was designated as the lung-cell donor and contributed lung, liver, and spleen for baseline virome screening (**Table S2**). NC01 also provided the uninoculated tissue control shown in Figure 3B. Individual animal characteristics and study contributions are summarized in Table S2.

Animals were maintained in the Tulane animal facility under controlled environmental conditions, a 12-h light/dark cycle, and ad libitum access to species-appropriate food and water. Welfare monitoring included general appearance, activity, intake, body weight, body temperature, and procedure-related adverse events. Procedures were approved by the Tulane University Institutional Animal Care and Use Committee and performed under institutional EBV biosafety requirements.

### Baseline Virome Screening of Chinese Tree Shrew Tissues

Lung, liver, and spleen from uninoculated animal NC07 (**Table S2**) were examined for detectable viral RNA as an organ-level baseline assessment. Total RNA was isolated with TRIzol and analyzed by rRNA-depleted RNA sequencing. Reads were screened against a composite reference containing the Chinese tree shrew TS3.0 genome and 746 mammalian viral genomes, including Tupaia herpesvirus strain 2, tree shrew papillomavirus 1, tree shrew papillomavirus 2, and tree shrew polyomavirus 1 (5, 21, 22). Alignments were generated with STAR and the reported default options (5, 23). Candidate viral alignments were inspected in IGV (24).

### Virus production and Raji-cell titration

Akata GFP-EBV, a widely used infectious recombinant derived from the Akata EBV strain, was used throughout this study. The recombinant carries a GFP reporter cassette, enabling identification of infected cells while preserving major latency-associated genomic regions, including the BART region and EBNA2 locus. Akata GFP-EBV producer cells were induced with anti-human IgG at 10 µg/mL for 72 h. Culture supernatants were harvested, clarified by centrifugation, passed through 0.45-µm filters, aliquoted, and stored at -80°C. Serial dilutions were applied to Raji cells, and GFP-positive recipient cells were used to express infectious activity as green Raji units per milliliter (GRU/mL).

### In vivo inoculation and longitudinal sampling

Six Chinese tree shrews were inoculated intravenously through the tail vein with 290 µL of Akata GFP-EBV preparation per animal. Based on the infectious titer of 8.93 × 10⁵ GRU/mL determined by Raji-cell titration, the administered dose was approximately 2.6 × 10⁵ GRU per animal. Blood and oral-swab specimens from TS01-TS05 were collected at nonuniform, animal-specific intervals over approximately two years, with additional collections beyond two years in TS02 and TS05. Sampling times are shown as days post-inoculation in the individual profiles (**Fig. 2B**). Animal characteristics and study contributions are summarized in Table S2. This observation period was not a common terminal endpoint for all animals.

TS06 underwent a separate blood and oral-swab monitoring series through day 174 post-inoculation and was euthanized for tissue collection at that endpoint. Whole blood was collected into anticoagulant-containing tubes, and oral-swab specimens were placed in phosphate-buffered saline (PBS) before DNA extraction.

### Uninoculated control monitoring

Blood and oral-swab specimens from six uninoculated tree shrews (NC01-NC06) were collected intermittently during the control observation period and tested for EBV DNA. The number and spacing of collections varied among animals. Animal characteristics are summarized in Table S2, and the individual blood and oral profiles are presented in Fig. S1. Time in Fig. S1 is expressed as days from the first collection for each animal.

### EBV DNA qPCR in blood and oral specimens

DNA was extracted with the DNeasy Blood and Tissue Kit (Qiagen). EBV genomic targets were amplified on a Bio-Rad real-time PCR instrument, with technical triplicates. Absolute EBV DNA copy numbers were determined using a standard curve generated from serial dilutions of EBV Akata BACmid DNA and expressed as copies/mL of blood or copies/swab. Nondetected specimens are indicated as ND. For display on logarithmic axes, ND specimens were placed at a fixed plotting position that does not represent a measured viral concentration or the assay detection limit. Comparisons across time use the same sample-processing and calibration convention.

### Tissue collection and EBER RNAscope

Liver, lung, and spleen sections were examined from two inoculated animals sampled at different stages: TS06 at day 174 post-inoculation and TS01, a member of the longitudinal cohort, at day 720 (approximately two years). Corresponding tissues from the uninoculated animal NC01 served as controls. Five-micrometer formalin-fixed, paraffin-embedded sections were prepared for hematoxylin and eosin staining and EBV-encoded RNA (EBER) RNAscope. The EBER probe (ACDBio, Cat# 310271) was used according to the manufacturer’s instructions. EBV-positive and EBV-negative controls were included in each run. Digital slides were acquired with a Motic EasyScan Pro 6 scanner (Motic, Hong Kong, China), and representative fields were captured at 40× magnification as previously described (4). EBER findings were recorded separately for each donor and organ and are summarized in Table S3.

### Receptor sequence analysis

Human CR2/CD21 (NP_001006659.1, 1,092 amino acids), Chinese tree shrew CR2/CD21 (XP_014447439.1, 1,091 amino acids), human EphA2 (NP_004422.2, 976 amino acids), and tree shrew EphA2 (TSDBTID.7937, 976 amino acids) were compared using pairwise protein alignments analyzed with custom Python scripts. Amino-acid identity was calculated as the number of identical residue pairs divided by the total number of alignment columns, including columns containing a gap, multiplied by 100. Residue numbering refers to the respective full-length precursor sequences. Positive BLOSUM62 substitution scores were used to annotate nonidentical residues in the alignment displays and were not counted as amino-acid identities. The CR2 N-terminal SCR1-SCR2 region was mapped using the human fragment represented by PDB entry 1LY2. The displayed comparison spans human residues 22-149 and corresponding tree shrew residues 22-150. Human EphA2 domain intervals were taken from UniProt P29317 and projected onto the tree shrew sequence through the protein alignment. Regional comparisons included the extracellular region (human residues 24-537), ligand-binding domain (28-206), and first and second fibronectin type-III domains (328-432 and 438-529). Targeted alignments are shown in Figs. 3C and 4B. Full-length CR2 and EphA2 alignments are provided in Figs. S2 and S3, respectively. Table S4 summarizes regional coordinates, identical-residue counts, alignment lengths, gap columns, and identity percentages.

### Establishment of lung-derived cultures and EBV-positive derivatives

Primary cultures were generated from lung tissue of an uninoculated tree shrew (designated donor NC07, **Table S2**) and immortalized independently by retroviral expression of hTERT or T antigen, followed by puromycin selection. EBV-positive derivatives were generated by cell-to-cell exposure to induced Akata GFP-EBV producer cells and maintained under G418 selection at 100 µg/mL. Established cultures were maintained in RPMI 1640 with 10% fetal bovine serum.

### RT-qPCR analysis

cDNA was synthesized from total RNA using iScript gDNA Clear cDNA Synthesis Kit (BioRad, Cat# 172-5035). qPCR analysis was performed using iQ SYBR Green Supermix (Bio-Rad, Cat# 170-8882) on a Bio-Rad CFX96 instrument as follows: 1 μl of cDNA and 1 μl of 10 μM primers were mixed with 10 μl of SYBR green supermix and 8 μl nuclease-free H2O to a 20 μl reaction volume. Polymerase was activated and cDNA was denatured at 95°C for 5 minutes. cDNA was then amplified for 40 cycles with 15 s denaturation at 95°C, 60 s annealing/extension and plate reading at 60°C. Melting curve analysis was performed at temperatures from 60°C to 90°C with 0.5°C increment per 5 s. Technical triplicates were averaged for each biological sample before calculating group means. Primer sequences are listed in Table S1. Relative transcript expression was calculated using the 2^−ΔΔCt^ method after normalization to GAPDH, with the corresponding EBV-negative parental culture as the calibrator within each immortalization background. For viral targets without detectable amplification in the parental controls after 40 cycles, a Ct value of 40 was assigned for calculation of relative expression. Values calculated using Ct = 40 are relative to an assigned control baseline, not a measured control transcript abundance.

### Transfection

Transient transfection experiments were performed by using a modified version of the calcium phosphate precipitation procedure (25). 1 x 10^6^ cells were plated onto 100-mm-diameter tissue culture dishes. The following day, the medium was replaced with 8 ml of fresh supplemented DMEM. Four hours later, 0.5 ml of 1x HEPES-buffered saline (0.5% HEPES, 0.8% NaCl, 0.1% dextrose, 0.01% anhydrous Na2HPO4, 0.37% KCl [pH 7.10]) was mixed with a total of 30 µg of plasmid DNA (effector plasmids and carrier DNA [pUHD10] were added to make a total of 30 µg). 30 µl of 2.5 M CaCl2 was added, and samples were mixed immediately. Precipitates were allowed to form at room temperature for 20 minutes before being added dropwise to cells. Cells were incubated at 37°C with 5% CO2 for 16 h before the medium was replaced with 10 ml of fresh supplemented DMEM. Cells were incubated for another 24 h before harvesting.

### Sodium butyrate treatment

EBV-positive hTERT- and TAg-immortalized tree shrew lung-derived cultures were treated with 3 mM sodium butyrate (NaB). Cells were harvested after 48 h for Western blot analysis of EBV lytic protein expression. Samples for EBV DNA quantification and donor-culture supernatants for the Raji infectivity assay were collected after 72 h of treatment and processed as described below.

### Western blot analysis

The cells were pelleted at desired time point and washed with 1X PBS and then the cells were immediately suspended in 100 μl of 1× Laemmli sample buffer and heated for 15 minutes at 95°C before electrophoresis. Protein concentrations of the whole-cell extracts were measured with the Bio-Rad protein assay kit according to the manufacturer’s instructions. Equal amounts of protein were subjected to SDS-polyacrylamide gel electrophoresis and transferred to PVDF membranes (Whatman). The blots were blocked for 60 minutes in Tris-buffered saline containing 5% nonfat powdered milk and 1% FBS and then incubated with the primary antibody (in blocking buffer) overnight at 4°C. The blots were washed three times with 1× TBST (140 mM NaCl, 3 mM KCl, 25 mM Tris-HCl [pH 7.4], 0.1% Tween 20) (each wash was carried out for approximately 15 minutes). To detect the chemiluminescence signals, the blots were then incubated with horseradish peroxidase-conjugated secondary antibody (Bio-Rad) in blocking buffer for 1 h at room temperature. Blots were washed as described above and analyzed with an enhanced chemiluminescence detection system (Perkin-Elmer) according to the manufacturer’s recommendations. Signals were acquired using a Bio-Rad ChemiDoc imaging system or by exposure to Fuji Super RX films (Fujifilm) followed by film scanning. To detect infrared fluorescence signals, an Odyssey infrared imaging system (Li-Cor Biosciences) was used with a secondary antibody (IRDye 680RD goat anti-rabbit antibody, Li-Cor, Cat# 925-68071). Immunoblot figure panels were assembled from the original image files using uniform brightness/contrast or levels adjustments applied to the entire blot image or panel. No selective enhancement, erasure, cloning, masking, background filling, or local manipulation was performed.

The following primary antibodies were used for Western blot analysis: anti-Zta MAb (Santa Cruz, Cat# SC-53904), anti-BMRF1 MAb (Santa Cruz, Cat# SC-58121), anti-VCA/p18 (ThermoFisher, Cat# PA1-73003), anti-EBNA1 (Santa Cruz, Cat# SC-81581), anti-CD21 (Santa Cruz, Cat# SC-13135), anti-EphA2 (Cell Signaling Technology, Cat# 6997T), anti-GAPDH (Santa Cruz, Cat# SC-47724), and anti-alpha/beta-Tubulin antibody (Cell Signaling Technology, Cat# 2148S).

### Analysis of DNase-resistant EBV DNA

Cell-free EBV DNA in culture supernatants was assessed using DNase-resistant DNA measurements. Cells were pelleted (300 × g, 10 minutes, 4°C), and supernatants were clarified (2,000 × g, 10 minutes, 4°C) and passed through 0.45-µm syringe filters to remove residual cells and debris. Clarified supernatants were aliquoted and subjected to DNase treatment or mock treatment in parallel. For DNase digestion, Turbo DNase I (Thermo Fisher) was added to a final concentration of 20 U/mL in the supplied buffer (final 5 mM MgCl₂, 1 mM CaCl₂) and incubated 1 h at 37°C. Reactions were stopped by adding EDTA to 10 mM and heating 75°C for 10 minutes, followed by lysis with proteinase K (200 µg/mL) and 0.5% SDS for 30 minutes at 55°C. Viral DNA was extracted from 200 - 500 µL of treated or mock-treated supernatant using the QIAamp MinElute Virus Spin Kit (Qiagen) and eluted in 50 µL of nuclease-free water. EBV DNA was quantified by qPCR targeting EBV genomic noncoding sequences using validated primer sets. Primer sequences are listed in Table S1. Absolute copy numbers were interpolated from plasmid standards.

### Infectious EBV production assay

Culture supernatants were collected from EBV-positive lung-derived donor cultures treated with 3 mM NaB for 72 h and from matched control cultures, then clarified by centrifugation to remove cells and debris. Supernatants were then filtered through a 0.45 μm membrane. Viral particles were concentrated from clarified supernatants by polyethylene glycol (PEG) precipitation according to the laboratory protocol. The resulting pellet was resuspended in RPMI medium and stored at -80°C until use. Concentrated viral preparations were added to Raji cells. After incubation, GFP-positive Raji cells were quantified by flow cytometry. Infectious EBV production was expressed as the percentage of GFP-positive Raji cells. Vehicle-treated donor cultures served as negative controls.

### RNA isolation, library preparation, and sequencing

Total RNA from the six lung-derived cultures was isolated using TRIzol and treated with DNase. RNA concentration and integrity were assessed with an Agilent Bioanalyzer. Strand-specific, rRNA-depleted total-RNA libraries were sequenced as 100-bp paired-end reads on a DNBSEQ-500 platform. Approximately 48 million read pairs per sample were retained after filtering. The six sequenced libraries comprised one parental comparator and two independently established infected derivatives in each immortalization background, not six replicates of one condition.

### RNA-seq quality control and reference preparation

Read quality was assessed with FastQC v0.12.1, adapters and low-quality sequence were removed with fastp v1.1.0, and reports were summarized with MultiQC v1.35. Reads were analyzed against the premerged Chinese tree shrew TS3.0 reference (22, 26, 27) and Akata EBV genome (18). The viral contig was designated chrEBV_Akata_inverted, with promoter-associated Cp-EBNA and Qp-EBNA1 transcript models retained. Human-ortholog gene symbols in the tree shrew annotation supported the lineage and pathway analyses. Salmon inference and an independent featureCounts comparison supported an inward-facing, reverse-stranded configuration for all six epithelial libraries (ISR, featureCounts -s 2). Transcript identifiers were retained consistently across the annotation and quantification output. Transcript sequences for the combined reference were extracted with gffread.

### Genome alignment, transcript quantification, and gene counting

Reads were aligned to the combined genome with STAR v2.7.10b, retaining coordinate-sorted BAM files and splice-junction tables. BAM indexing and alignment-quality summaries were generated with SAMtools. Strand assignment was applied during quantification and feature counting rather than described as a restriction on genomic alignment. Primary host counts were generated with featureCounts v2.1.1 using paired-read counting, reverse-strand assignment (-s 2), exclusion of multimapping reads, and exclusion of fragments assigned ambiguously to more than one gene. The opposite setting (-s 1) was examined as a quality-control comparison. Transcript abundance was estimated with Salmon using ISR, sequence-bias and GC-bias corrections, and the combined transcriptome. Transcript estimates were summarized with tximport. Fractional viral counts were kept separate from the integer counts used for host differential expression.

### Viral expression, coverage, and promoter-associated junctions

EBV abundance was expressed as TPM and summarized both including and excluding EBER1/2. Conservative viral counts were accompanied by fractional-overlap estimates to assess the influence of overlapping annotations. Coverage was normalized per million mapped alignments and displayed in 500-bp bins. Breadth was calculated against the 171-kb Akata EBV coordinate system. Cp-and Qp-associated EBNA1 transcript estimates remained separate, and unique splice-junction reads were collapsed by genomic coordinate before comparison with the expected promoter-associated junctions. Latent-associated, lytic-associated, BART-region, and other viral modules were retained. Per-base depth and normalized bedGraph tracks were generated with SAMtools and BEDTools v2.31.1, respectively.

### Host differential expression and lineage assessment

Host counts were analyzed in R v4.4.2 with DESeq2 using ∼ background + ebv_status, controlling for hTERT versus T-antigen background. Differential expression was evaluated with Wald tests from negative-binomial generalized linear models, with Benjamini-Hochberg adjustment of P values across tested host genes. Log2 fold changes were shrunk with apeglm. Primary robust genes required adjusted P ≤ 0.05, absolute shrunken log2 fold change ≥ 1, concordant direction across backgrounds, agreement in at least three of four infected-derivative-versus-parent comparisons, and direction stability in at least three of four leave-one-infected-line-out analyses. Exploratory candidates displayed in the cross-background comparison and ranked supplementary heatmap required adjusted P ≤ 0.10, absolute shrunken log2 fold change ≥ 1, concordant direction across backgrounds, and agreement in at least three of four derivative-versus-parent comparisons, without the leave-one-line-out requirement. The high-confidence subset additionally required adjusted P ≤ 0.01, absolute shrunken log2 fold change ≥ 1.5, and complete line-level and leave-one-line-out directional agreement. For lineage-marker assessment, expression was displayed as log2(TPM + 1), using TPM ≥ 1 as a descriptive detection threshold. Module scores were mean log2(TPM + 1) across available markers.

### Pathway enrichment and transcriptome visualization

Preranked enrichment was performed with fgsea using the MSigDB Hallmark, Reactome, and Gene Ontology Biological Process collections obtained through msigdbr. Tree shrew genes were mapped to human ortholog symbols. Where multiple genes mapped to the same symbol, the gene with the largest absolute ranking statistic was retained for that ranked list. The blocked-model DESeq2 Wald statistic was the primary ranking measure, with background-specific log2 fold-change rankings considered descriptively. Enrichment was calculated using fgseaMultilevel for gene sets containing 10-500 represented genes, and P values were adjusted by the Benjamini-Hochberg procedure within each collection and ranking analysis. Prioritization required blocked-model FDR ≤ 0.10 and concordant enrichment direction across backgrounds. The displayed Hallmark sets met FDR ≤ 0.05. PCA and Spearman sample correlations used variance-stabilized counts. Host differential-expression heatmaps are described below. Other heatmaps used the transformations stated in their legends. Data handling and visualization used dplyr, tidyr, tibble, readr, ggplot2, ggrepel, and pheatmap. The RNA-seq workflow also used Python v3.12.2 scripts.

### Host-expression heatmaps

The heatmap (**Fig. 7C**) displays 24 high-confidence genes selected to illustrate the examples discussed in the Results, including all seven targets assessed by RT-qPCR. The eight increased and 16 decreased genes were grouped by direction and ordered by increasing adjusted P value within each group. The supplementary heatmap (**Fig. S4**) displays the 50 exploratory candidates with the lowest adjusted P values, with ties resolved by decreasing absolute shrunken log2 fold change. Rows and columns were hierarchically clustered. Both heatmaps used the same six RNA-seq libraries. For each gene, variance-stabilized counts were centered by subtracting the mean across all six cultures and divided by the sample standard deviation across those cultures to obtain row z scores.

### Statistical analysis

Non-transcriptomic comparisons were analyzed in GraphPad Prism 9. Technical qPCR triplicates were averaged within each biological sample and were not treated as independent biological replicates. For the cellular qPCR comparisons indicated in the figure legends, measurements were analyzed on the log2 scale before two-sided t tests against the corresponding control. For relative expression calculated by the comparative Ct method, log2 relative expression equals −ΔΔCt. Data are presented as mean ± SEM unless otherwise stated. Animal ages are reported as mean ± sample SD, referenced to inoculation for TS01-TS06, first blood collection for NC01-NC06, or tissue collection for NC07. Longitudinal animal measurements and qualitative tissue findings were summarized descriptively. RNA-seq gene-level and pathway analyses used the multiple-testing adjustments and selection thresholds specified above. For the targeted comparisons, P < 0.05 was the nominal significance threshold. The reported asterisk thresholds are * P < 0.05, ** P < 0.01, *** P < 0.001, and **** P < 0.0001.

### Database accession number

The RNA-seq sequence data have been deposited in the NIH Sequence Read Archive (SRA) database under accession number PRJNA1555165.

## RESULTS

### Establishment of a Chinese tree shrew colony for integrated in vivo and in vitro EBV studies

To provide a sustained source of animals and species-matched cells for EBV research, we established a Chinese tree shrew colony at Tulane University (**Fig. 1A**). The study involved 13 individual animals. The experimental framework integrated longitudinal blood and oral monitoring, tissue EBER localization in TS06 and the long-term animal TS01, and a lung-derived cellular system for viral transcription and lytic-induction studies (**Fig. 1B**).

**Figure 1.**
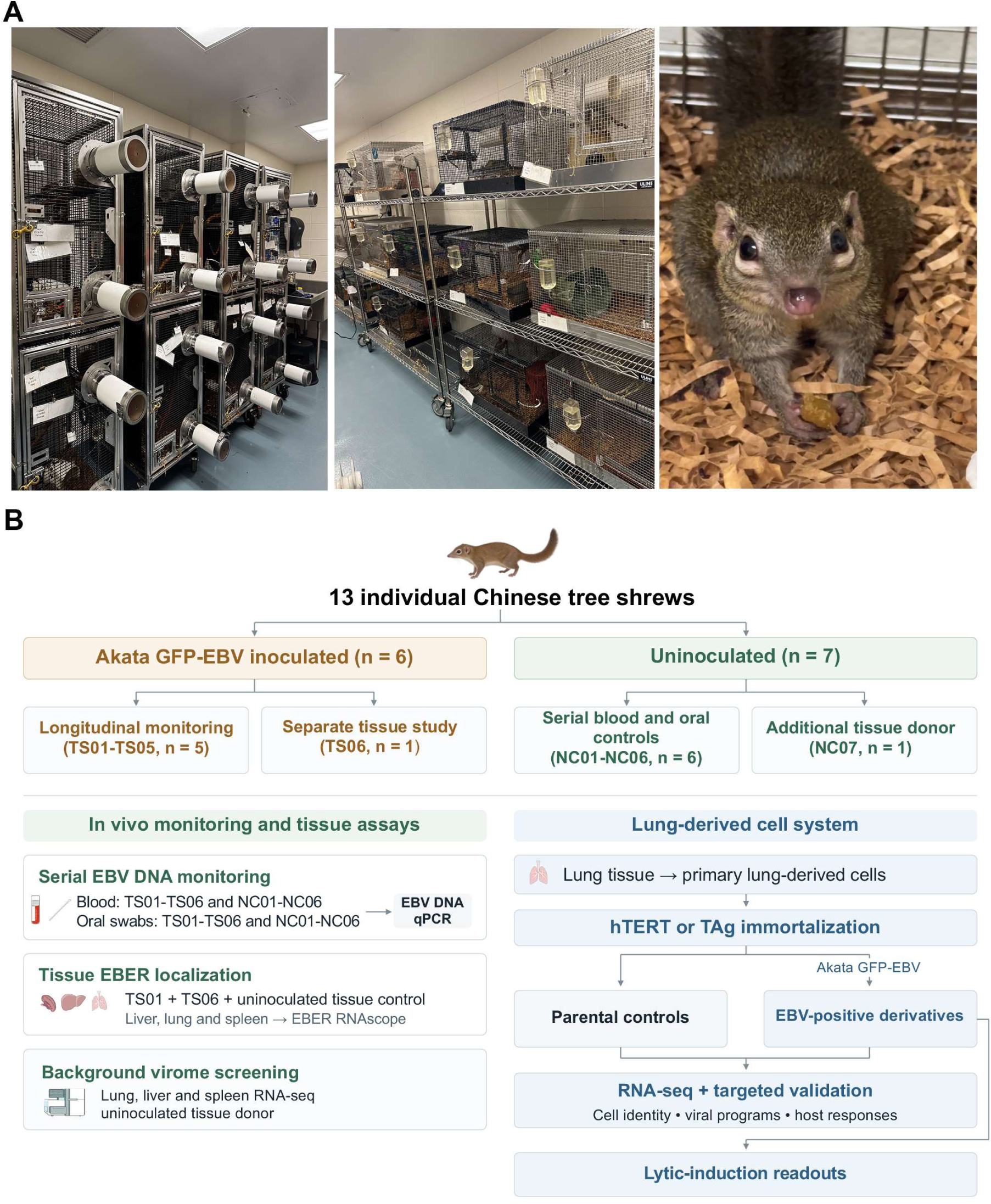
Establishment of a Chinese tree shrew colony and an integrated animal-cell platform for EBV research. (A) Representative photographs of the tree shrew housing facility and a colony animal. (B) Experimental framework involving 13 individual tree shrews: six Akata GFP-EBV-inoculated animals (TS01-TS06) and seven uninoculated animals (NC01-NC07). TS01-TS05 formed the longitudinal cohort, with sampling beyond two years in TS02 and TS05. TS01 also contributed tissues near two years. TS06 underwent serial blood and oral monitoring followed by terminal tissue collection at day 174. NC01-NC06 provided serial blood and oral controls, and NC07 was the designated lung-cell donor and baseline-virome donor (Table S2). Liver, lung, and spleen were examined by EBER RNAscope from TS06, TS01, and the uninoculated control NC01. Background virome screening used lung, liver, and spleen from NC07. The cell arm included hTERT- and T-antigen-immortalized parental cultures and EBV-positive derivatives evaluated by RNA-seq, targeted assays, and lytic-induction studies. Individual study contributions are summarized in Table S2, serial uninoculated-control measurements in Fig. S1, and tissue EBER findings in Table S3.

To characterize the baseline virologic background of the colony and assess the potential presence of endogenous or preexisting viral infections that could complicate subsequent EBV studies, we performed RNA-seq-based virome screening of lung, liver, and spleen tissues from uninoculated animal NC07 (**Table S2**). Sequencing data were analyzed against a comprehensive host-viral reference containing 746 mammalian viral genomes. No viral sequences meeting the established detection criteria were identified for the surveyed human-infecting viruses or four previously reported tree shrew viruses, including *Tupaia herpesvirus* strain 2, tree shrew papillomaviruses 1 and 2, and tree shrew polyomavirus 1. These findings demonstrated the absence of detectable RNA from the surveyed viruses in the examined tissues, providing a favorable baseline virologic profile for subsequent experimental EBV infection studies.

### Longitudinal monitoring reveals intermittent EBV DNA detection extending beyond two years

To establish the in vivo EBV infection model, we first quantified the infectious activity of the Akata GFP-EBV preparation using GFP-based Raji-cell titration. Serial dilutions were applied to Raji indicator cells, yielding an infectious titer of 8.93 × 10⁵ green Raji units (GRU)/mL (**Fig. 2A**). Six Chinese tree shrews were inoculated intravenously with 290 µL of this preparation per animal, corresponding to approximately 2.6 × 10⁵ GRU. Five animals (TS01-TS05) underwent longitudinal monitoring, while TS06 was designated for a separate monitoring and tissue-analysis study.

**Figure 2.**
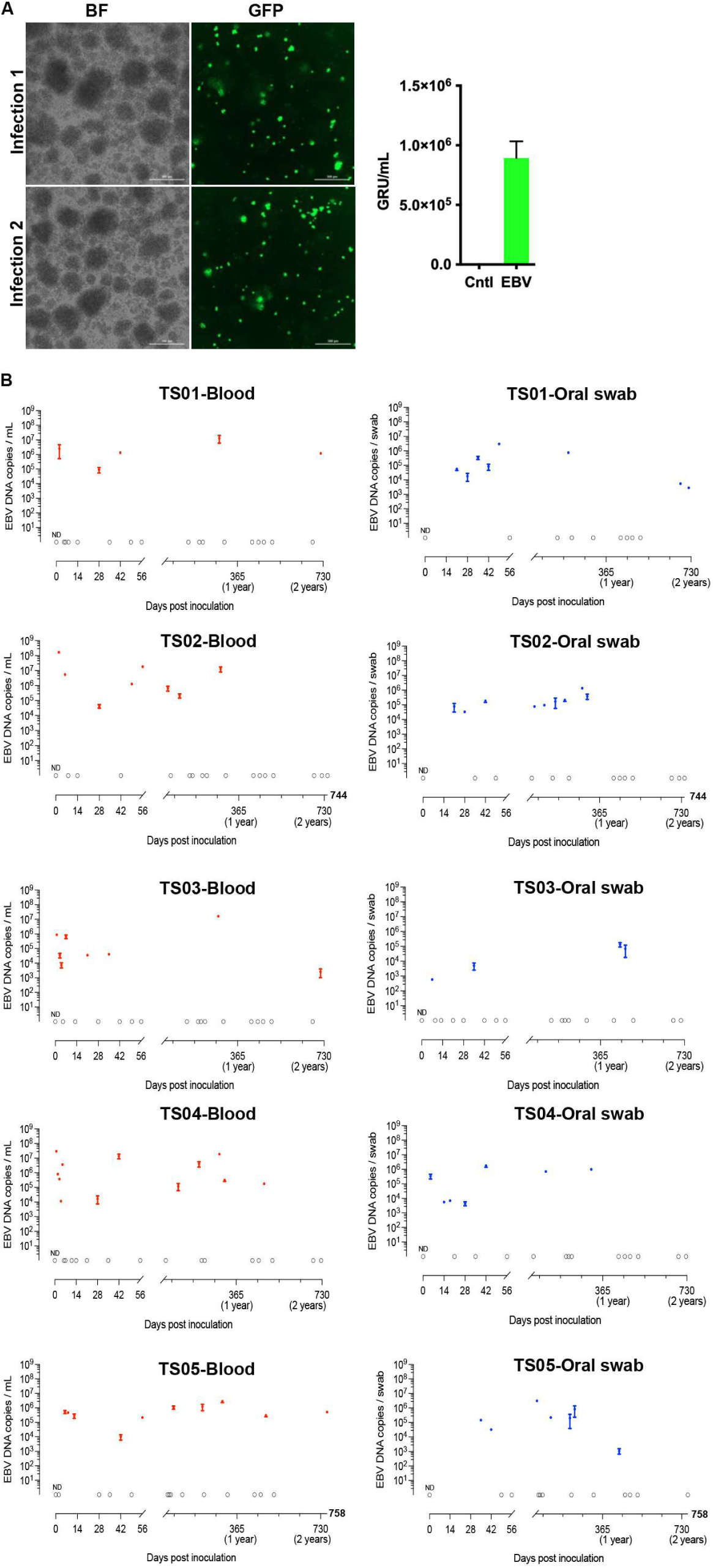
Longitudinal monitoring reveals intermittent blood and oral EBV DNA detection, including blood positivity beyond two years. (A) GFP-based Raji-cell titration of Akata GFP-EBV, with representative bright-field (BF) and GFP images and a stock titer of 8.93 × 10⁵ green Raji units (GRU)/mL. Scale bars, 300 μm. (B) EBV DNA measurements in blood (left) and oral-swab specimens (right) from TS01-TS05. Each animal is displayed separately, with observations plotted at their actual days post-inoculation. Sampling intervals varied among animals. TS02 and TS05 contributed specimens beyond two years, and a post-two-year blood specimen from TS05 was EBV DNA-positive. Values are expressed as copies/mL of blood or copies/swab on logarithmic axes. Filled symbols indicate detected measurements. Open symbols at ND indicate tested specimens without reportable detection. Their fixed vertical position is a display convention, not a measured concentration or detection limit. The early and subsequent observations are shown on separate x-axis segments. The 730-day tick marks a two-year reference and is not a common final sampling date. Serial uninoculated-control measurements are shown in Fig. S1.

To assess prolonged EBV detectability, we monitored blood and oral-swab specimens from TS01-TS05 at nonuniform intervals (**Fig. 2B**). Sampling extended to approximately two years in the longitudinal cohort and beyond two years in TS02 and TS05. All five animals had detectable EBV DNA in both specimen types during the initial monitoring period. Positive and nondetected measurements occurred intermittently, with individual variation in the timing and magnitude of detection.

Importantly, EBV DNA was detected in a blood specimen from TS05 collected more than two years after inoculation (**Fig. 2B**). Selected blood and oral specimens were also positive during late follow-up (**Fig. 2B**). These observations extend documented viral detectability beyond the initial post-inoculation period and support long-term EBV persistence. Furthermore, EBV DNA was not detected in any tested blood or oral-swab specimen from the six uninoculated controls (NC01-NC06) during their collection period, providing biological negative-control observations for both specimen types (**Fig. S1 and Table S2**).

### Tissue EBER localization supports multiorgan infection and long-term EBV persistence

To complement the longitudinal viral DNA measurements with cellular localization in tissues, we examined liver, lung, and spleen from two inoculated animals sampled at different stages. TS06 underwent serial blood and oral-swab monitoring through day 174, followed by terminal tissue collection (**Fig. 3A**). EBER RNAscope detected cell-associated signals in all three organs from TS06, whereas corresponding sections from the uninoculated control NC01 lacked specific staining (**Fig. 3B**). In the liver, some EBER-positive cells displayed morphology consistent with hepatocytes. In the TS06 lung, signals were associated with cells along alveolar septa, suggesting possible epithelial involvement. Importantly, EBER signals were also detected in liver and spleen sections from TS01 near two years after inoculation, but not in the examined lung sections (**Fig. 3B and Table S3**). Together, these observations demonstrate multiorgan localization of EBV-associated RNA. The late liver and spleen findings in TS01 complement the serial blood and oral measurements from the same animal and provide tissue-level evidence supporting long-term EBV persistence.

**Figure 3.**
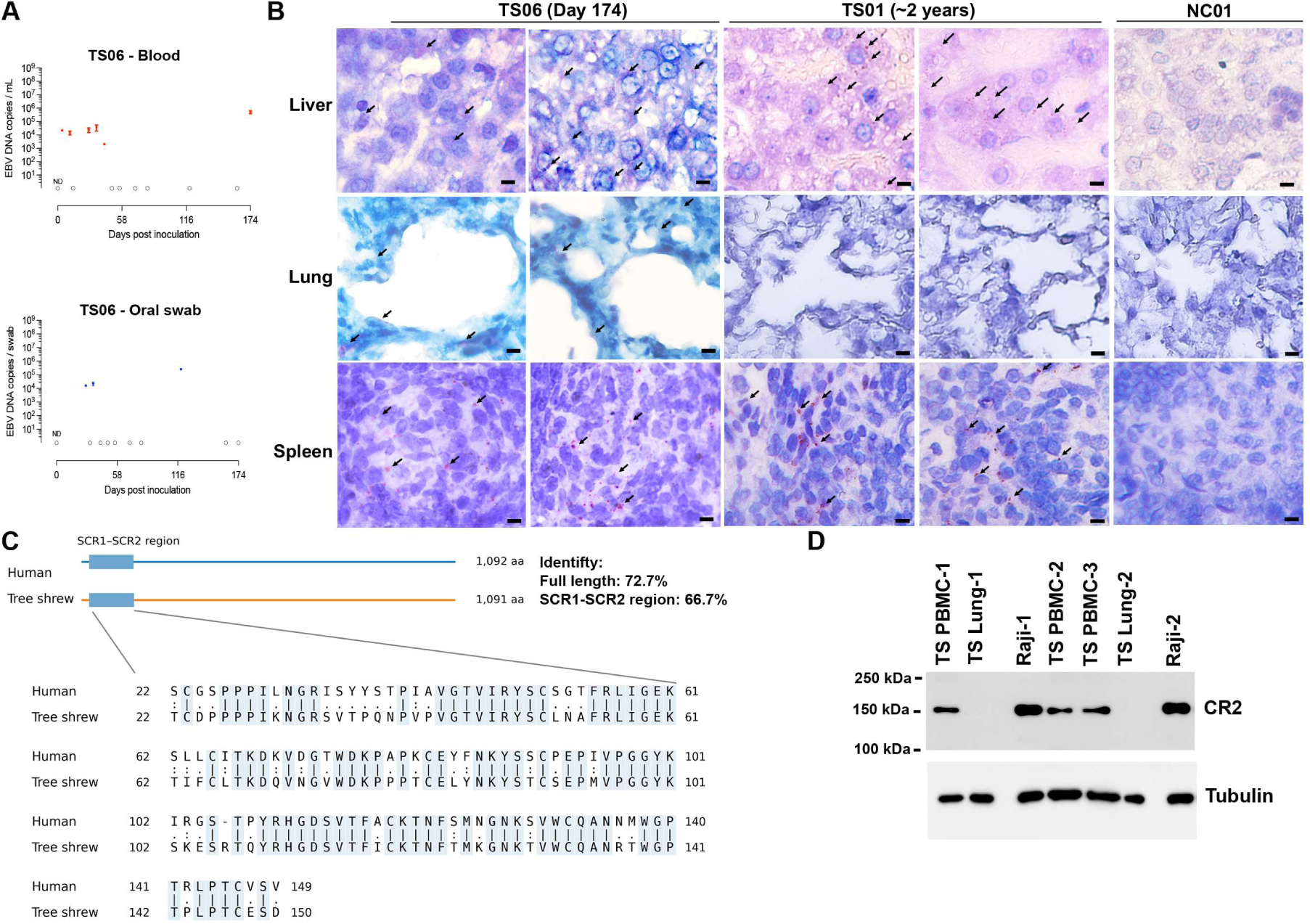
Tissue EBER localization at earlier and late collection points and CD21/CR2 characterization. (A) Blood (upper) and oral-swab (lower) EBV DNA measurements from TS06 through terminal collection at day 174. Filled symbols indicate detected DNA. Open ND symbols indicate tested nondetects. Their vertical position does not represent a measured concentration or detection limit. Units are copies/mL of blood and copies/swab. (B) EBER RNAscope of liver, lung, and spleen from TS06 (day 174), TS01 (approximately two years), and uninoculated control NC01. Two representative fields per organ are shown for each inoculated animal and one for NC01. Arrows indicate cell-associated EBER signals. Scale bars, 10 μm. Some signal-bearing cells in the liver exhibit hepatocyte-like morphology, while positive cells in the TS06 lung are situated along alveolar septa. EBER signals were detected in all three examined organs from TS06 and in liver and spleen from TS01, but not in the examined TS01 lung or NC01 sections. The donor-by-organ findings are summarized in Table S3. (C) Targeted CR2 SCR1-SCR2 alignment: human NP_001006659.1, residues 22-149, and tree shrew XP_014447439.1, residues 22-150. Identity is 66.7% (86/129 columns, including gaps). Shading/vertical bars indicate identity. Colons, positive BLOSUM62 substitutions. Dots, other substitutions. Dashes, gaps. Coordinates refer to precursor proteins. The full-length alignment is in Fig. S2, with calculations in Table S4. (D) Western blot analysis of CD21/CR2 expression in three tree shrew peripheral blood mononuclear cell (PBMC) preparations and two lung tissue lysates, with human Raji B cells included as a positive control. CD21-immunoreactive bands were detected at approximately 150 kDa in all three tree shrew PBMC preparations and in the Raji positive control, consistent with the expected molecular weight of CR2. No comparable bands were detected in the lung tissue lysates under the conditions examined. Tubulin served as the loading control.

To characterize a receptor context relevant to tree shrew EBV susceptibility, we examined complement receptor 2 (CR2/CD21), a major attachment receptor for EBV in human B cells. Earlier work reported conservation of selected gp350-binding residues and CR2 expression in tree shrews (12). Comparison of the human and tree shrew reference proteins identified 72.7% full-length amino-acid identity (795 identical residues across 1,094 alignment columns). The 1LY2-mapped N-terminal SCR1-SCR2 region, spanning human residues 22-149 and tree shrew residues 22-150, showed 66.7% identity (86/129 alignment columns), with one gap-containing column in the targeted alignment shown in Fig. 3C. The full-length alignment is provided in Fig. S2, and identity calculations are summarized in Table S4. To complement these sequence comparisons, we examined CD21 immunoreactivity by Western blotting with an anti-human CD21 antibody. A band at approximately 150 kDa was detected in each of the three displayed tree shrew PBMC preparations and in the positive-control lanes, whereas neither of the two whole-lung preparations showed a comparable band under the conditions examined (**Fig. 3D**). These complementary sequence and immunoblot observations further characterize CD21/CR2 in the tree shrew model.

### Establishment of a tree shrew lung-derived cell system supporting EBV infection and EphA2 expression

To complement our *in vivo* EBV infection model, we sought to establish a renewable, species-matched cell culture system for investigating EBV infection, latency, and lytic reactivation. Although Chinese tree shrews have been used to investigate EBV infection *in vivo*, a well-characterized lung-derived cellular system for studying EBV infection and viral gene regulation in this species has been lacking. To address this gap, we established primary lung-derived cultures from the uninoculated donor NC07 and immortalized them independently using human telomerase reverse transcriptase (hTERT) or T antigen. We subsequently established stable Akata GFP-EBV-positive derivatives in both immortalization backgrounds. Representative bright-field and fluorescence images demonstrated GFP expression in the infected cultures (**Fig. 4A**), establishing a tractable cellular platform for further characterization of EBV infection.

**Figure 4.**
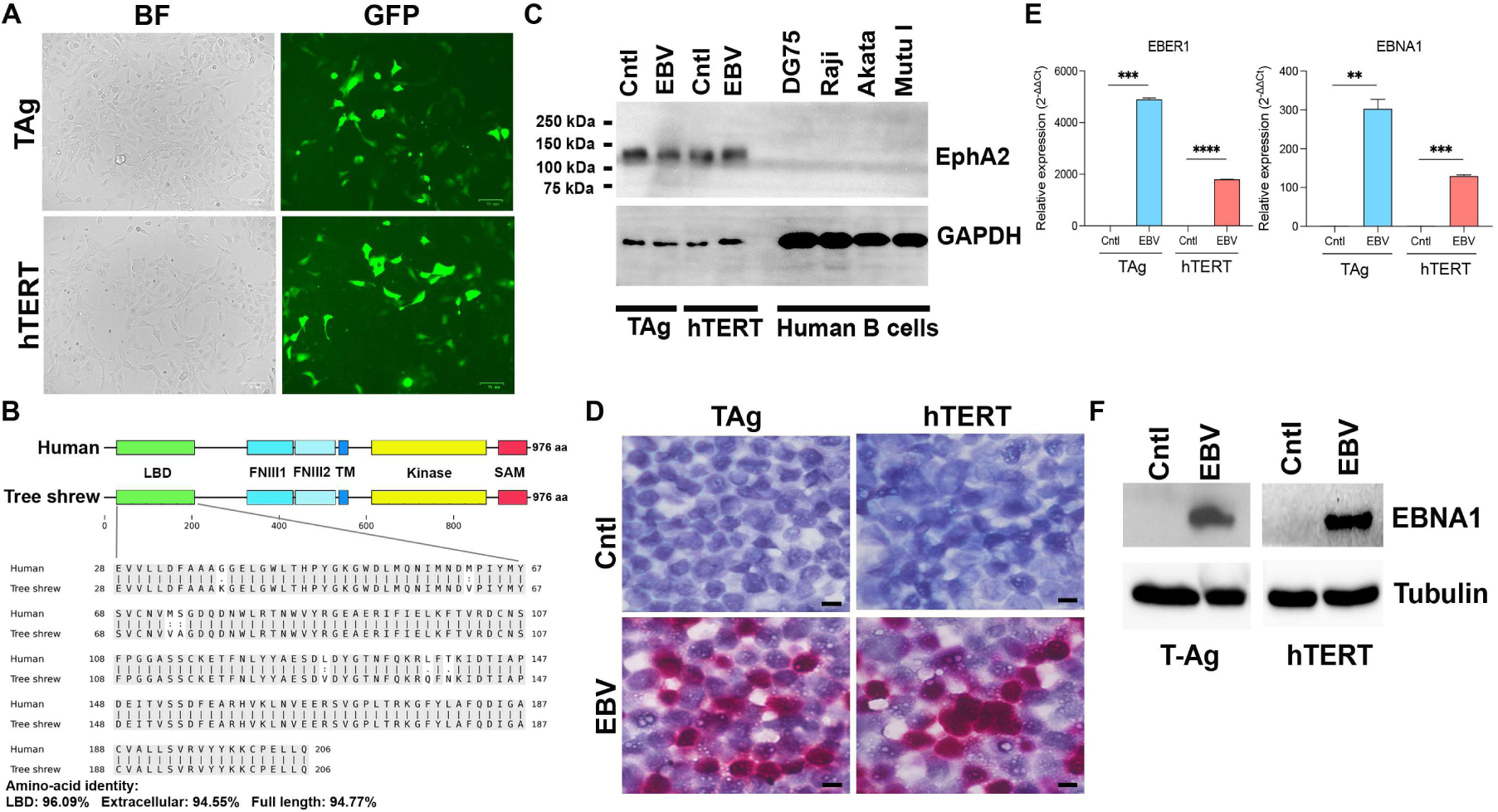
Establishment of EphA2-expressing tree shrew lung-derived cultures supporting maintained EBV infection. (A) Bright-field (BF) and GFP images of EBV-positive T-antigen (TAg)- and hTERT-immortalized cultures. Scale bars, 71 μm. (B) Protein-domain overview and targeted EphA2 ligand-binding-domain alignment: human NP_004422.2 and tree shrew TSDBTID.7937, precursor residues 28-206. Regional identity is 96.09% (172/179 columns), without gaps. Shading/vertical bars indicate identity. Colons, positive BLOSUM62 substitutions. Dots, other substitutions. Human domain boundaries follow UniProt P29317. The full-length alignment is in Fig. S3 and regional identity calculations are in Table S4. LBD, ligand-binding domain. FNIII1/2, fibronectin type-III domains. TM, transmembrane region. SAM, sterile alpha motif. (C) EphA2 immunoblot analysis of EBV-negative parental and EBV-positive tree shrew lung-derived cultures in both immortalization backgrounds. Human B-cell lines DG75, Raji, Akata, and Mutu I were included as negative controls for EphA2 expression. No detectable EphA2 bands were observed in these B-cell lysates under the conditions examined. GAPDH served as the loading control. (D) EBER RNAscope in parental (Cntl) and infected cultures. Scale bars, 10 μm. (E) EBER1 and EBNA1 RT-qPCR, displayed as relative expression calculated using the 2^−ΔΔCt^ method. Nondetected viral targets in parental controls were assigned Ct = 40 for calculation. These values are relative to an assigned, not a measured, viral-expression baseline. Gray denotes parental controls, blue the TAg infected comparison, and red the hTERT infected comparison. Data are reported as means ± SEM from ≥2 biological preparations, each assayed in technical triplicate. Comparisons used two-sided t tests on log2-transformed relative-expression values versus the indicated control. *, P < 0.05, **, P < 0.01, ***, P < 0.001, and ****, P < 0.0001. (F) EBNA1 immunoblots with tubulin loading controls. TAg is shown on the left and hTERT on the right.

Because EphA2 is an established entry receptor for EBV in human epithelial cells (19, 20), we next evaluated conservation of its reference protein and expression in the tree shrew lung-derived cultures. The human and tree shrew EphA2 sequences each contained 976 amino acids and shared 94.77% full-length identity (925/976 residues) without alignment gaps. Identity was 94.55% in the extracellular region (residues 24-537) and 96.09% in the ligand-binding domain (28-206), with corresponding coordinates in both sequences. The targeted ligand-binding-domain alignment is shown in Fig. 4B, the full-length alignment in Fig. S3, and regional identity calculations in Table S4. The first and second fibronectin type-III domains showed 92.38% and 94.57% identity, respectively. These comparisons demonstrate that the high overall identity extends to the extracellular receptor regions. Western blotting with an anti-human EphA2 antibody detected prominent immunoreactivity in both parental and EBV-positive cultures from the hTERT and T-antigen backgrounds (**Fig. 4C**). Together, these results demonstrate substantial EphA2 sequence conservation and complementary immunoreactivity in the lung-derived cell system.

To further characterize EBV infection in these newly established cell lines, we examined the expression of viral latency-associated RNA and protein markers. EBER RNAscope demonstrated strong EBER signals in EBV-positive derivatives (**Fig. 4D**). Consistent with these findings, RT-qPCR detected EBER1 and EBNA1 transcripts in the infected cultures from both immortalization backgrounds (**Fig. 4E**). Western blot analysis further confirmed EBNA1 protein expression in EBV-positive cells but not in their corresponding parental controls (**Fig. 4F**). Together, these complementary findings demonstrate the maintenance of EBV-associated viral RNA and protein expression in tree shrew lung-derived cells and establish a species-matched cellular platform for investigating EBV latency and lytic reactivation.

### RNA-seq reveals reproducible EBV-positive cell states and a lung-derived epithelial/mesothelial-like phenotype

Having established stable EBV-positive tree shrew lung-derived cultures, we next sought to characterize their cellular identity and determine whether independently generated infected derivatives exhibited reproducible transcriptional profiles. Although the cultures displayed epithelial-like morphology and expressed EphA2, their molecular identity had not been comprehensively characterized. Furthermore, because both immortalization and EBV infection can influence cellular gene expression, transcriptomic profiling was important for distinguishing the effects of different immortalization backgrounds and establishing a foundation for subsequent analyses of EBV-associated transcriptional changes. To address these questions, we performed strand-specific, rRNA-depleted RNA sequencing of six cell cultures, comprising two EBV-negative parental lines and four independently established EBV-positive derivatives generated using hTERT or T-antigen immortalization.

Approximately 48 million paired-end fragments were retained per library. Reverse-stranded feature counting assigned 36.5 - 38.6% of the reported alignment total, compared with 7.7 - 7.9% using the opposite-strand convention, confirming the expected library orientation. Principal component analysis (PCA) distinguished the two immortalization backgrounds and separated EBV-negative parental cultures from their corresponding EBV-positive derivatives. PC1 and PC2 accounted for 53.8% and 25.6% of the observed transcriptomic variance, respectively (**Fig. 5A**). Notably, independently established EBV-positive derivatives clustered closely within each immortalization background and exhibited high transcriptome-wide Spearman correlations (**Fig. 5B**), demonstrating reproducible transcriptional profiles among the infected cultures.

**Figure 5.**
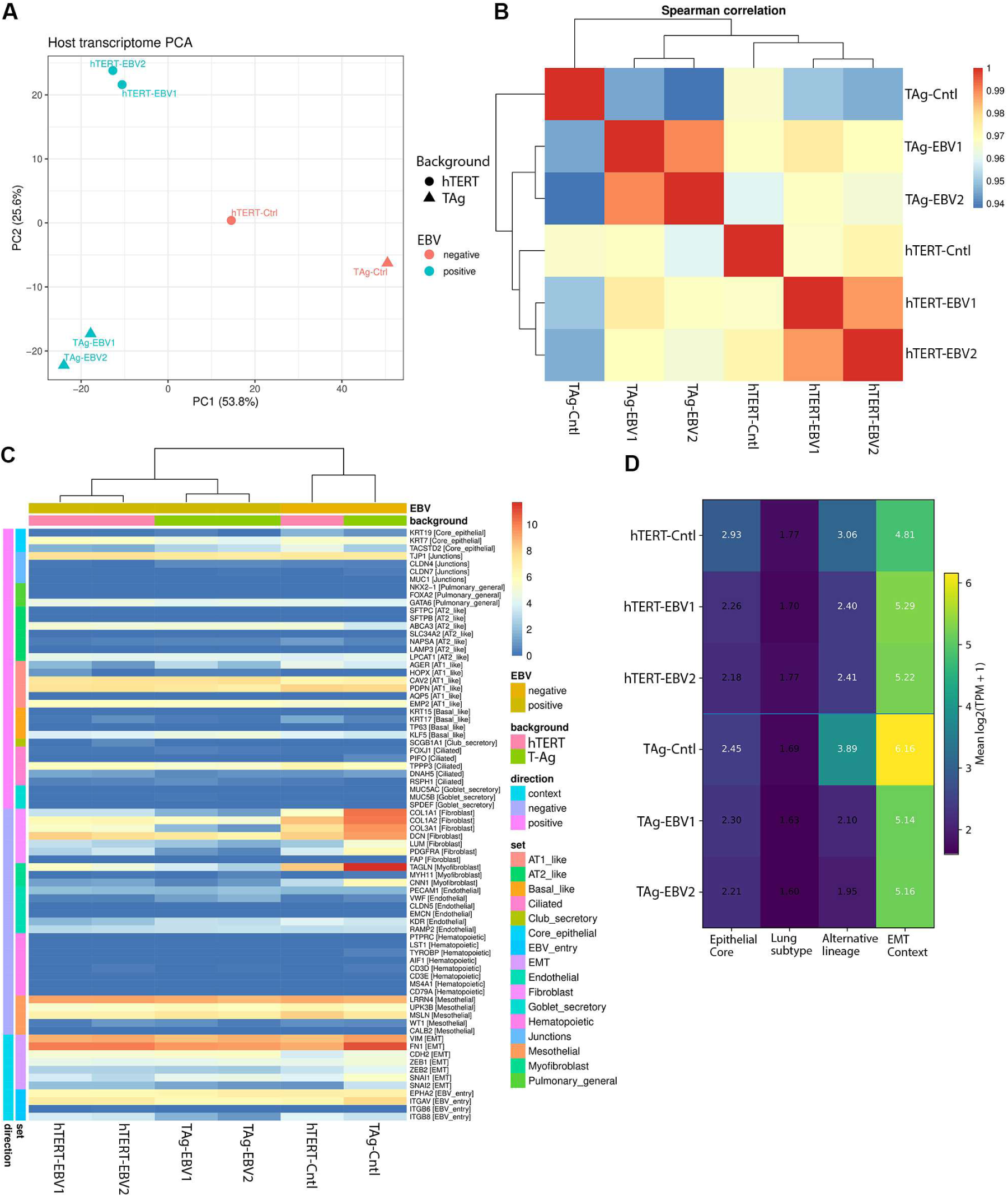
RNA-seq reveals reproducible transcriptional profiles and an epithelial/mesothelial-like phenotype in tree shrew lung-derived cells. Six tree shrew lung-derived cell cultures were analyzed, comprising two immortalization backgrounds. The hTERT group included an EBV-negative parental line (hTERT-Cntl) and two independently established EBV-positive derivatives (hTERT-EBV1 and hTERT-EBV2). Similarly, the T-antigen (TAg) group included TAg-Cntl, TAg-EBV1, and TAg-EBV2. (A) Principal component analysis (PCA) of host RNA-seq profiles. PC1 and PC2 explain 53.8% and 25.6% of the variance, respectively. Shapes indicate immortalization background, and colors indicate EBV infection status. (B) Spearman correlation heatmap with hierarchical clustering, showing similarities in host gene expression among the six cultures. (C) Heatmap of selected epithelial, pulmonary, mesothelial, mesenchymal, hematopoietic, and EBV entry-associated genes. Expression is presented as log₂(TPM + 1). Annotation bars indicate EBV status, immortalization background, marker categories, and marker interpretation. The direction annotation denotes lineage-marker interpretation rather than differential-expression direction. (D) Mean log₂(TPM + 1) expression scores for epithelial-core, pulmonary-subtype, alternative-lineage, and epithelial–mesenchymal transition (EMT) gene modules. Together, these analyses demonstrate reproducible transcriptional profiles among EBV-positive derivatives and identify a lung-derived epithelial/mesothelial-like phenotype with partial mesenchymal features.

To further define the cellular identity of these lung-derived cultures, we examined the expression of established epithelial, pulmonary, mesothelial, and mesenchymal lineage markers. All six cultures expressed epithelial and cell junction-associated genes, including KRT7, TACSTD2, TJP1, and GATA6, together with selected pulmonary-associated genes and the EBV epithelial entry-related factors EPHA2 and ITGAV (**Fig. 5C**). However, markers associated with differentiated alveolar and airway epithelial cells, including NKX2-1, SFTPC, SFTPB, SLC34A2, AQP5, and SCGB1A1, were absent or expressed at low levels. Interestingly, the cultures also exhibited prominent expression of mesothelial-associated genes, including LRRN4, MSLN, and UPK3B, together with mesenchymal-associated genes such as FN1, VIM, CDH2, and ZEB1. In contrast, hematopoietic lineage markers were negligible. Consistent with these observations, gene-module analyses revealed a mixed transcriptional profile encompassing epithelial, mesothelial, and mesenchymal features (**Fig. 5D**).

Together, these findings establish the molecular characteristics of our newly developed tree shrew lung-derived cell system, demonstrating reproducible transcriptional profiles among independently established EBV-positive derivatives and identifying an epithelial/mesothelial-like phenotype with partial mesenchymal features. This characterization provides an important foundation for subsequent investigations of EBV latency, lytic gene expression, and virus-associated host transcriptional remodeling.

### Viral profiling identifies Qp-associated EBNA1 transcription alongside latent-associated and spontaneous lytic expression

To determine how EBV gene expression is organized after infection is established in tree shrew lung-derived cells, we examined viral transcript abundance, promoter-associated EBNA1 models and splice junctions, and latent-associated and lytic genes. This analysis was designed to extend characterization beyond viral-marker detection by identifying features of transcriptional regulation in the maintained cultures.

Quantifiable EBV transcription was detected exclusively in the four EBV-positive derivatives. Conservative feature counting assigned 3,191 - 6,961 EBV fragments per infected library, whereas no viral fragments were assigned to either parental library. Total EBV transcript abundance ranged from 1,255 to 1,785 summed TPM across the infected lines. Notably, 524 - 1,090 TPM remained after excluding the highly abundant nonpolyadenylated EBER1 and EBER2 transcripts (**Fig. 6A**), demonstrating that EBV transcription extended well beyond EBER expression.

**Figure 6.**
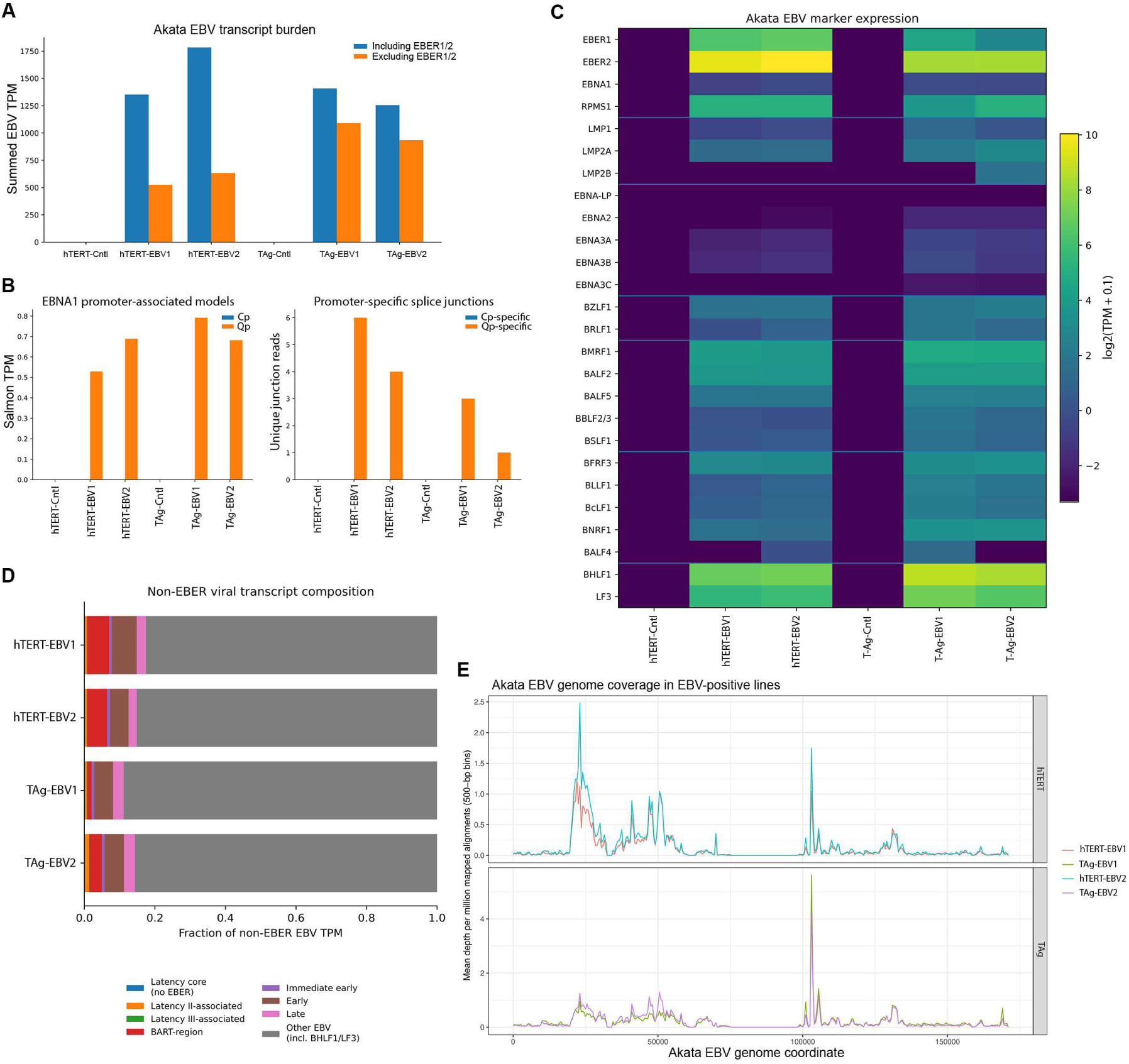
Viral profiling identifies Qp-associated EBNA1 transcription alongside latent-associated and spontaneous lytic expression. (A) Summed viral TPM, including and excluding EBER1/2, in the six cultures defined in Figure 5. (B) Cp- and Qp-associated EBNA1 transcript estimates and unique reads supporting the displayed promoter-specific junctions. Low-count Qp junctions provide qualitative promoter-associated evidence. (C) Latent-associated and lytic-associated transcripts displayed as log₂(TPM + 0.1). (D) Composition of non-EBER viral TPM in the four infected derivatives. The other-viral-transcript category includes BHLF1 and LF3. (E) Akata EBV RNA-seq coverage normalized per million mapped alignments and averaged in 500-bp bins, separated by immortalization background. Parental cultures lacked quantifiable viral transcription. The hTERT and TAg coverage panels use different y-axis scales.

We next examined the viral latency-associated transcriptional program. All four infected derivatives expressed EBER1/2, EBNA1, LMP1, LMP2A, and RPMS1/BART-region transcripts. The RNA-seq estimates also included low-abundance signals for selected Latency III-associated genes, including EBNA2 and EBNA3-family members (**Fig. 6C**). To characterize EBNA1 transcriptional regulation, we analyzed promoter-associated transcripts and splice-junction evidence. Qp-associated EBNA1 transcripts were detected in the infected cultures and supported by a small number of Qp-specific splice-junction reads, whereas the corresponding Cp-specific junctions were not detected (**Fig. 6B**). These observations support Qp-associated EBNA1 transcription within a latent-associated expression profile that also includes selected broader EBNA transcripts.

Alongside latent-associated expression, we detected transcription of multiple EBV lytic genes in the absence of experimental induction. These included the immediate-early genes BZLF1 and BRLF1, early genes BMRF1, BALF2, and BALF5, and late genes BFRF3, BLLF1, and BcLF1 (**Fig. 6C**). BHLF1 and LF3 were also prominent among the non-EBER viral transcripts. Analysis of the non-EBER transcriptional composition showed contributions from multiple viral gene classes and was dominated by the other-viral-transcript category, which includes BHLF1 and LF3 (**Fig. 6D**). Consistent with this broad transcriptional activity, viral RNA-seq reads covered approximately 72 - 81% of the Akata EBV genome at a depth of at least one read, with reproducible but nonuniform coverage patterns (**Fig. 6E**). These findings establish that maintained latent-associated transcription coexists with spontaneous lytic gene expression in the bulk cultures.

Collectively, these findings identify Qp-associated EBNA1 transcription within a viral-expression profile that includes selected EBNA2 and EBNA3-family transcripts and spontaneous lytic genes. The promoter-associated and gene-level measurements provide a regulatory context for testing whether the maintained infection retains functional lytic competence.

### Cross-background host profiles identify E2F/MYC-associated enrichment and reduced immune-related transcription

Having characterized the EBV transcriptional programs maintained in tree shrew lung-derived cells, we next investigated whether EBV-positive status was associated with coordinated changes in host gene expression. EBV infection can profoundly influence cellular proliferation, metabolism, and immune signaling, contributing to the establishment and maintenance of infected cell populations. However, the host transcriptional landscape associated with EBV infection in tree shrew cells remains largely unexplored. To address this question, we performed differential gene expression and pathway enrichment analyses comparing EBV-positive derivatives with their corresponding uninfected parental cultures. Importantly, analyses incorporated both hTERT- and T-antigen-immortalized backgrounds to identify reproducible transcriptional changes across independently established cellular systems.

Among 15,501 host genes examined, 684 met our primary robustness criteria, which incorporated statistical significance, effect size, and directional consistency across immortalization backgrounds (**Fig. 7A**). The descriptive cross-background plot highlights the exploratory candidates (**Fig. 7B**). A more stringent analysis identified 395 high-confidence differentially expressed genes, including 54 increased and 341 decreased genes in EBV-positive cells. Examples of decreased genes in the high-confidence subset included FMOD, SULF1, SERPING1, PCDH19, SYNPO2, CHI3L1, VCAM1, TNFAIP2, PDGFRB, COL6A1, TNC, CD248, CD74, ITGA11, IGFBP2, and NCAM1, many of which are associated with extracellular matrix organization, cell adhesion, and immune-related functions. Examples of increased genes included NRG1, ATP8A1, TMC7, TIAM1, PARM1, PLA2G3, PRKCZ, and TP73.

**Figure 7.**
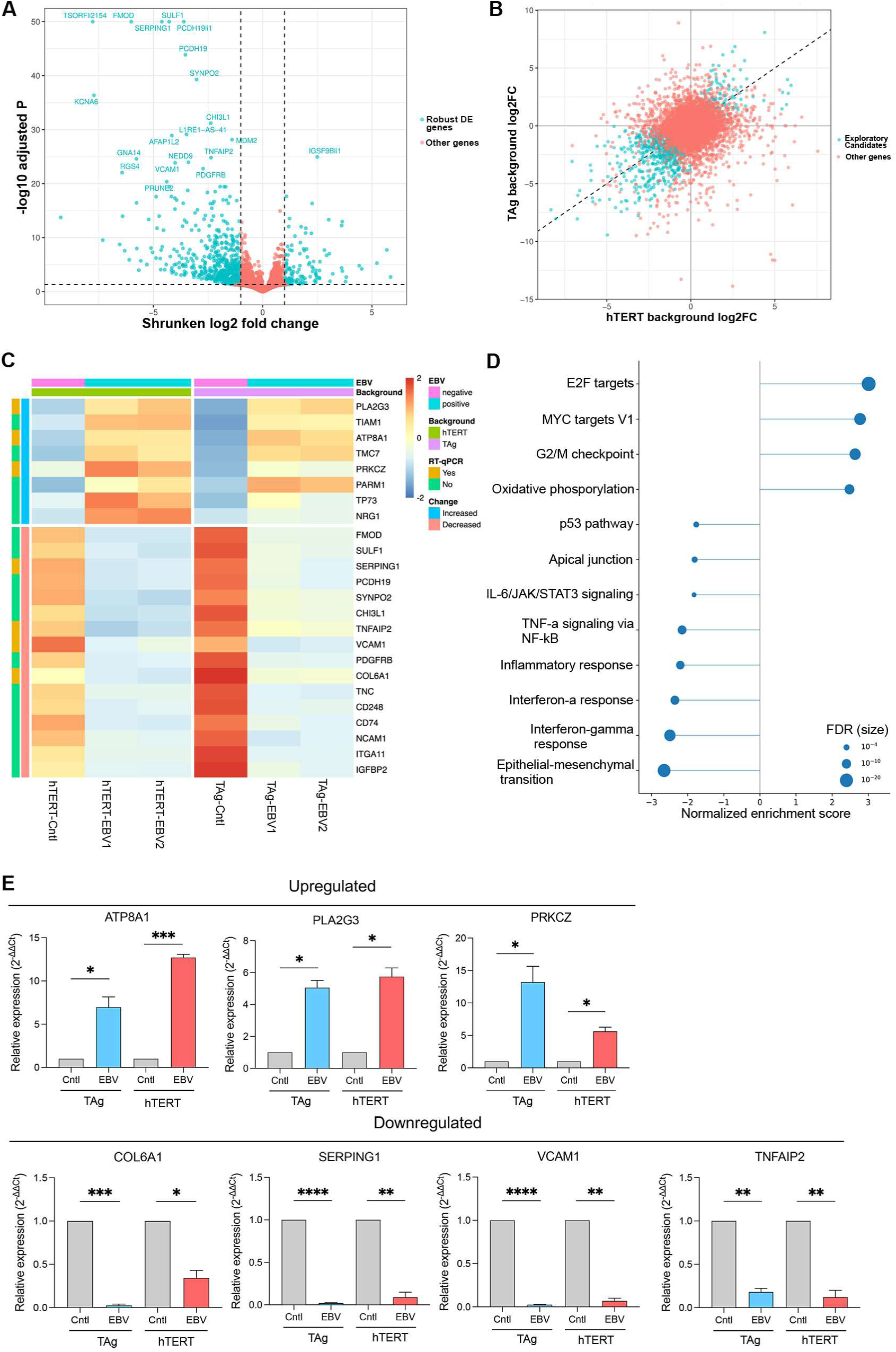
Host transcriptional remodeling across immortalization backgrounds and RT-qPCR corroboration of selected genes. (A) Volcano plot from the blocked DESeq2 model (∼ background + ebv_status), showing shrunken log₂ fold change and −log₁₀ adjusted P value, capped at 50 for display. Cyan identifies primary robust differentially expressed (DE) genes and salmon identifies other genes. Dashed lines indicate adjusted P = 0.05 and log₂ fold change = ±1. (B) Descriptive log₂ fold changes in the hTERT and T-antigen backgrounds. Cyan identifies exploratory candidates, a different selection from panel A. Salmon identifies other genes. The dashed diagonal indicates equal fold changes. Selection criteria are defined in Methods. (C) Heatmap of 24 selected high-confidence host genes discussed in the Results, comprising eight increased and 16 decreased genes. Variance-stabilized counts are displayed as row z scores calculated within each gene across all six cultures. Genes are grouped by direction of change and ordered by increasing original adjusted P value within each group. The complementary ranked exploratory heatmap is shown in Fig. S4. (D) Selected Hallmark enrichment results. Positive normalized enrichment scores (NES) indicate enrichment toward the EBV-positive end of the ranked list. Negative scores indicate enrichment toward the parental end. Point area scales with −log₁₀(FDR). Larger points indicate lower FDR. (E) RT-qPCR assessment of increased ATP8A1, PLA2G3, and PRKCZ and decreased COL6A1, SERPING1, VCAM1, and TNFAIP2 relative to the corresponding parental cultures. Gray denotes controls, blue the T-antigen infected comparison, and red the hTERT infected comparison. All seven targets agree directionally with RNA-seq in both backgrounds. Data represent means ± SEM from ≥2 biological replicates, each assayed in technical triplicate. Comparisons used two-sided t tests on log2-transformed relative-expression values versus the indicated control. *, P < 0.05, **, P < 0.01, ***, P < 0.001, and ****, P < 0.0001.

These 24 selected high-confidence genes are displayed across the six cultures in Fig. 7C. The panel includes all seven host targets examined by RT-qPCR and illustrates concordant expression changes within both immortalization backgrounds. A complementary heatmap of the 50 exploratory candidates with the lowest adjusted P values is presented with hierarchical clustering in Fig. S4. Both panels visualize subsets of the same RNA-seq analysis.

To characterize the biological programs associated with these transcriptional changes, we next performed gene set enrichment analysis (GSEA). Hallmark pathway analysis revealed strong positive enrichment of E2F targets (normalized enrichment score [NES] = 3.01, FDR = 3.8 × 10⁻²³), MYC targets V1 (NES = 2.77, FDR = 2.0 × 10⁻¹⁶), G2/M checkpoint (NES = 2.64, FDR = 2.0 × 10⁻¹⁵), and oxidative phosphorylation (NES = 2.48, FDR = 5.4 × 10⁻¹²), suggesting increased expression of genes involved in cell-cycle progression, proliferation-associated processes, and cellular metabolism (**Fig. 7D**). Conversely, epithelial-mesenchymal transition (NES = -2.65, FDR = 1.1 × 10⁻¹⁹) and interferon-γ response (NES = -2.49, FDR = 9.1 × 10⁻¹⁶) were strongly negatively enriched. Additional negatively enriched pathways included interferon-α response, TNF/NF-κB signaling, inflammatory response, apical junction, p53 signaling, and IL-6/JAK/STAT3 signaling. Consistent with these findings, Reactome and Gene Ontology analyses provided complementary support for enrichment of proliferation-related and metabolic programs together with reduced extracellular matrix, cell adhesion, and immune-associated transcriptional programs. These coordinated results identify growth- and metabolism-associated expression signatures accompanied by reduced interferon- and inflammatory-response transcription in the EBV-positive cultures.

To assess selected RNA-seq findings using a complementary assay, we performed RT-qPCR analysis of seven high-confidence host genes included in Figure 7C. Consistent with the RNA-seq results, ATP8A1, PLA2G3, and PRKCZ were increased, whereas COL6A1, SERPING1, VCAM1, and TNFAIP2 were decreased in EBV-positive derivatives relative to their corresponding parental cultures (**Fig. 7E**). Importantly, all seven genes exhibited concordant directions of change in both hTERT- and T-antigen-immortalized backgrounds, although the magnitudes varied between the two systems. These findings corroborate the direction of selected EBV-associated host transcriptional changes using a complementary assay.

Together, the blocked analysis and cross-background agreement identify a coordinated EBV-associated host-expression state combining E2F/MYC-associated and metabolic enrichment with reduced matrix- and immune-related transcription. Directional agreement by RT-qPCR supports selected components of this state. This host profile provides a defined transcriptional context for the viral-expression and lytic-induction findings, rather than attributing the phenotype to a single immortalization background.

### Established EBV infection retains responsiveness to lytic induction and supports infectious output

To distinguish spontaneous lytic transcription from functional lytic competence, we asked whether established EBV-positive cultures could respond to induction with downstream viral protein expression and infectious output. We examined Zta/BZLF1 expression and sodium butyrate treatment in both immortalization backgrounds. These complementary interventions assessed responsiveness to a viral lytic regulator and to pharmacologic induction. Preparations from sodium-butyrate-treated culture supernatants were then evaluated in Raji recipients to test for infectious activity beyond intracellular viral-marker expression.

Western blot analysis demonstrated that both Zta expression and sodium butyrate treatment increased the expression of downstream EBV lytic proteins in the hTERT-and T-antigen-immortalized cell backgrounds (**Fig. 8A**). Specifically, induction was accompanied by increased detection of the early antigen EA-D/BMRF1 and the late viral capsid protein VCA p18. Importantly, the appearance of these downstream early and late proteins provides evidence that lytic induction extends beyond expression of the immediate-early transactivator Zta. These findings demonstrate that EBV-positive tree shrew lung-derived cultures retain the capacity to respond to both genetic and pharmacologic lytic stimuli and express viral proteins associated with successive stages of the lytic program.

**Figure 8.**
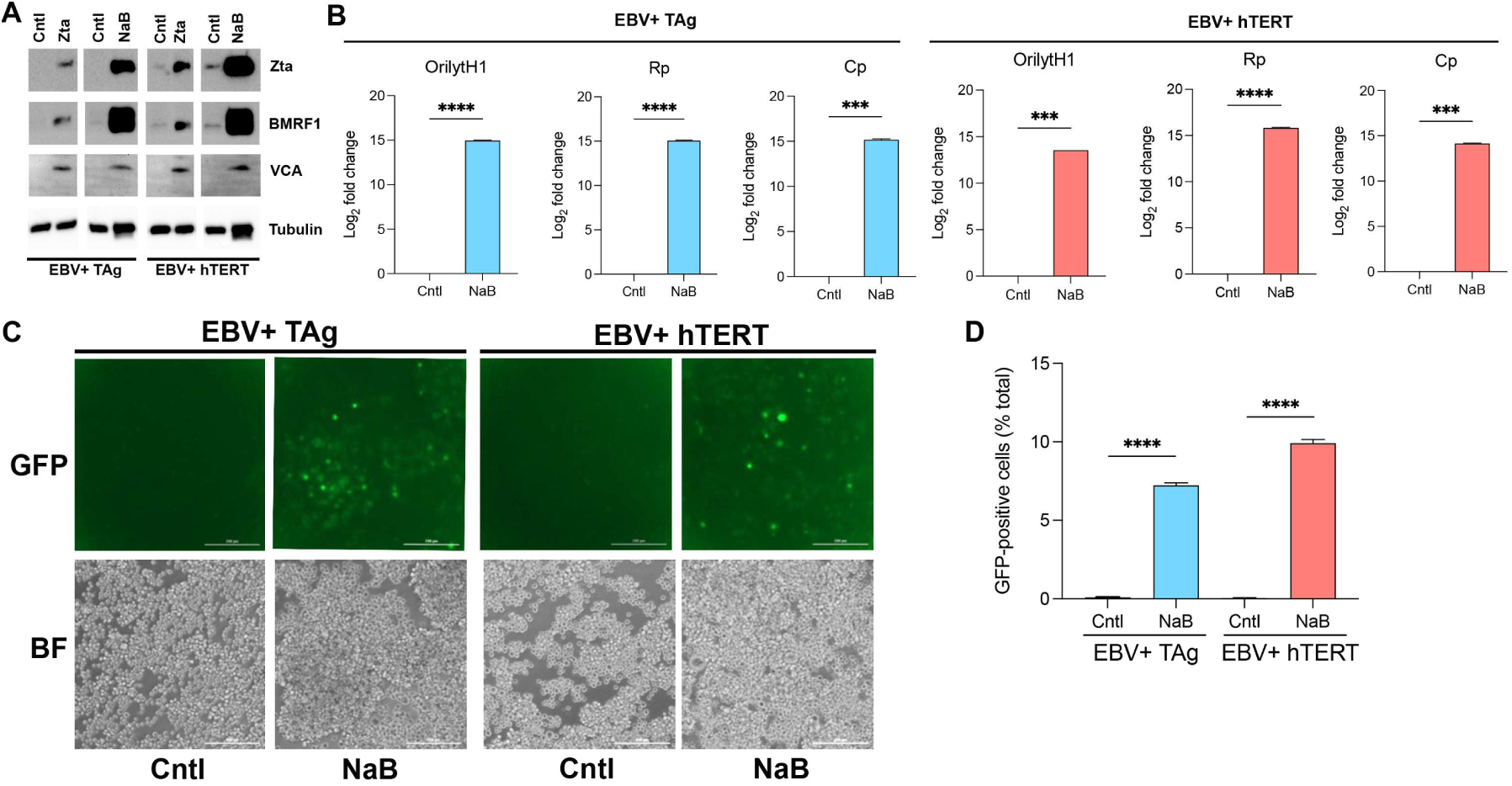
Established EBV infection retains inducible lytic protein expression and infectious output after sodium butyrate treatment. (A) Immunoblots of EBV-positive TAg and hTERT cultures following Zta expression or treatment with 3 mM sodium butyrate (NaB) for 48 h, with corresponding Cntl lanes. Zta, EA-D/BMRF1, and VCA/p18 represent immediate-early, early, and late viral protein readouts. Tubulin is the loading control. (B) DNase-resistant EBV DNA in cell-free culture supernatants collected from EBV-positive hTERT and TAg cultures after treatment with 3 mM NaB for 72 h or corresponding control treatment. Supernatants were treated with DNase before lysis and DNA extraction. Viral DNA was quantified by qPCR targeting the OriLytH1, Rp, and Cp genomic regions and is displayed as log₂ fold change. These measurements assess extracellular nuclease-protected viral DNA. Data are reported as means ± SEM from ≥2 biological preparations, each assayed in technical triplicate. Comparisons used two-sided t tests on log2-transformed DNA measurements versus the indicated control. *, P < 0.05, **, P < 0.01, ***, P < 0.001, and ****, P < 0.0001. (C) GFP and paired bright-field images of Raji recipients exposed to preparations recovered from supernatants of EBV-positive donor cultures treated with 3 mM NaB for 72 h or from corresponding control cultures. Column-group labels identify the donor-culture background. Scale bars, 200 μm. (D) Flow-cytometric quantification of GFP-positive Raji recipients, with approximately 7% positivity for induced TAg donors and 10% for induced hTERT donors.

To determine whether pharmacologic lytic induction promoted the release of EBV particles, we quantified DNase-resistant EBV DNA in cell-free culture supernatants collected after treatment with 3 mM sodium butyrate for 72 h. Supernatants were treated with DNase before lysis and DNA extraction, and viral DNA was quantified using qPCR assays targeting three distinct genomic regions, OriLytH1, Rp, and Cp. Sodium butyrate increased DNase-resistant EBV DNA at all three loci in both hTERT- and TAg-immortalized cultures (**Fig. 8B**). These concordant increases demonstrate accumulation of nuclease-protected extracellular EBV DNA, consistent with the release of genome-containing viral particles following lytic induction. This finding extends the evidence beyond intracellular viral protein expression to extracellular viral output.

To establish whether the released viral material was infectious, we next evaluated preparations recovered from sodium-butyrate-treated culture supernatants in Raji recipient cells. Concentrated preparations recovered from culture supernatants of sodium-butyrate-treated EBV-positive tree shrew cells were applied to Raji indicator cells, and infection-associated GFP expression was assessed by flow cytometry. Preparations from induced cultures generated substantial populations of GFP-positive Raji recipients, whereas the corresponding control preparations produced little detectable GFP signal (**Fig. 8C and D**). Approximately 7% and 10% of Raji recipient cells became GFP-positive following exposure to preparations recovered from the supernatants of induced T-antigen-and hTERT-derived cultures, respectively. These findings provide functional evidence consistent with the release of infectious Akata GFP-EBV from the induced tree shrew cells, extending the observations beyond viral gene expression and DNA detection.

Together, the protein, DNA, and recipient-cell measurements show that maintained EBV-associated infection in tree shrew cultures retains an inducible lytic program. Downstream early and late proteins respond to Zta expression or sodium butyrate, and the sodium-butyrate recipient assay provides functional evidence of infectious output. These findings extend the cellular platform beyond detection of infection-associated markers to functional assessment of viral activity after infection is established.

## DISCUSSION

This study establishes a complementary EBV platform that combines a small, immunocompetent host closely related to primates with regulatory and functional studies in species-matched lung-derived cultures. Blood EBV DNA detection beyond two years and EBER localization in liver and spleen near two years provide complementary evidence of long-term persistence. In the cultures, Qp-associated EBNA1 transcription, latent-associated and spontaneous lytic expression, and coordinated host-expression programs characterize the maintained infection. Downstream viral proteins respond to induction, and preparations from sodium-butyrate-treated cultures generate GFP-positive Raji recipients. The contribution therefore extends beyond demonstrating susceptibility to defining viral functions retained after infection has been established. These complementary animal and cellular systems expand the experimental scope of tree shrew EBV research alongside hematopoietic humanized mice (6, 7) and nonhuman-primate lymphocryptovirus models (8).

The tree shrew offers a useful combination of evolutionary proximity to primates and small-animal practicality. Comparative genomic studies support its close relationship to primates and provide increasingly detailed resources for investigating host pathways (22, 26, 27). Small body size and colony-based maintenance offer logistical advantages over larger nonhuman primates for serial sampling and extended infection studies (9, 10). Our colony and approximately two-year monitoring demonstrate the feasibility of this approach in the present setting. Unlike hematopoietic humanized mice, this model does not require human-cell engraftment and retains native immune, stromal, and epithelial compartments within one species. Direct use of Akata EBV also avoids exclusive reliance on a related simian lymphocryptovirus. The renewable cultures extend these advantages by providing a species-matched system for investigating maintained infection and inducible lytic activity. Together, these features connect the practical scale of a small animal with biological questions commonly addressed in primate infection models.

The longitudinal observations extend beyond the intervals reported in the cited earlier tree shrew studies, which followed infection for up to 21 weeks (11), 91 days (12), or 28 days (14). The positive blood specimen from TS05 establishes EBV DNA detectability beyond two years, rather than merely an extended collection schedule. EBER localization in liver and spleen of TS01 near two years adds a complementary tissue-level observation. These findings arose in different animals. Within TS01, the tissue results also connect late viral RNA detection with its own longitudinal specimen history. Together, the observations support persistence well beyond the initial post-inoculation phase. The absence of detectable EBV DNA in all tested blood and oral specimens from NC01-NC06 provides biological negative-control context for both specimen types (**Fig. S1**).

The use of Akata GFP-EBV provides an additional distinction from earlier studies employing deletion-bearing laboratory strains. The characterized Akata background (18) retains the BART region absent in B95-8 (15, 16) and the EBNA2 locus disrupted in P3HR-1 (17), while the GFP reporter permits visualization of infected cells and functional indicator-cell assays. Preservation of these genomic regions supports investigation of viral transcription in a more complete EBV background. In the present cultures, detection of RPMS1/BART-region RNA further demonstrates that this region contributes to the measured viral transcriptional program. Thus, strain selection and reporter-based readouts provide complementary advantages for linking molecular measurements to infection-associated cellular outcomes.

Establishment of an in-house colony also addresses a practical barrier to sustained tree shrew research by providing animals for longitudinal studies and tissue-derived cellular models. The baseline virome survey detected no RNA meeting the screening criteria for the surveyed human-infecting viruses or four specified tree shrew viruses in lung, liver, and spleen from uninoculated animal NC07 (**Table S2**). These findings provide a favorable baseline virologic profile for the examined tissues and complement the experimental characterization of the colony.

The tissue findings demonstrate EBV-associated RNA in multiple organs at the earlier collection and in liver and spleen near two years. No EBER signal was detected in the examined late TS01 lung sections. Because TS06 and TS01 are different animals and the measurements concern sampled sections, this comparison does not establish clearance from the lung or a time-dependent change in tissue tropism.

Morphology suggests hepatocyte involvement and possible pulmonary epithelial involvement, although definitive infected-cell lineages remain to be established. Receptor characterization provides additional cellular context. Earlier tree shrew work reported CR2 sequence conservation and expression (12). The present analysis identifies 72.7% full-length CR2 identity and 66.7% identity in the mapped SCR1-SCR2 region, alongside 94.77% full-length EphA2 identity and 96.09% identity in its ligand-binding domain. Targeted alignments are shown in Figs. 3C and 4B, with full-length alignments in Figs. S2 and S3 and identity calculations in Table S4. CD21 immunoreactivity in PBMCs and EphA2 immunoreactivity in lung-derived cultures complement these sequence comparisons. EphA2 conservation is relevant to its established role in human epithelial EBV entry (19, 20), although functional dependence on either receptor in tree shrew cells remains to be established.

A central resource developed here is the renewable lung-derived culture system. Stable EBV-positive derivatives were established independently in hTERT- and T-antigen-immortalized backgrounds, providing two settings in which viral and host phenotypes could be compared with corresponding parental cultures.

Transcriptomic characterization identifies a coherent epithelial/mesothelial-like phenotype with partial mesenchymal features, rather than assigning identity solely from tissue origin or morphology. Maintained EBER expression, EBNA1 RNA and protein, and reproducible viral transcription across independently established derivatives support the utility of these cultures for studying persistent EBV-associated cell states. The combination of defined lineage features, parental comparators, and two immortalization backgrounds makes the system an experimentally accessible complement to the animal model.

Viral profiling provides a regulatory layer to the cellular characterization. Qp-associated EBNA1 transcript models and supporting splice-junction reads identify a promoter-associated component of viral expression in the tree shrew cellular environment. Expression of EBER1/2, EBNA1, LMP1, LMP2A, and RPMS1 includes features of the restricted program described in epithelial EBV latency (28). Low-abundance RNA-seq estimates also support expression of selected Latency III-associated genes, including EBNA2 and EBNA3-family members. Immediate-early, early, and late transcripts reveal a spontaneous lytic component. These bulk observations describe coexisting transcriptional features rather than a uniform canonical latency state or exclusive promoter dependence. The combined profile places the induction experiments in a defined viral-expression context and provides a basis for investigating how maintained infection remains responsive to lytic regulation.

The host findings connect this viral-expression profile with a coordinated cellular state. The 684 robust EBV-associated expression differences, including a 395-gene high-confidence subset, were prioritized using concordance across immortalization backgrounds and infected derivatives. E2F- and MYC-associated gene sets, G2/M checkpoint, and oxidative phosphorylation were positively enriched, whereas matrix-, interferon-, and inflammatory-response programs were negatively enriched. The main and supplementary heatmaps illustrate selected components of this expression state using different gene-selection rules (**Figs. 7C and S4**). RT-qPCR corroborated seven selected targets in both backgrounds, providing complementary assay-level support for those genes. This combination provides a mechanistically relevant context for maintained infection and lytic inducibility. Our previous work showed that c-Myc and E2F1 can inhibit Zta-mediated transactivation in human epithelial-cell systems (25). Guo et al. subsequently identified MYC as a suppressor of EBV lytic reactivation in Burkitt lymphoma cells and linked this activity to regulation of contacts between oriLyt and the BZLF1 promoter (29). More recently, Biswas et al. showed that E2F1 represses BZLF1 transcription and positively regulates c-Myc in EBV-positive B-cell models (30). Together, these studies provide a mechanistic rationale for investigating whether the E2F/MYC-associated host state observed here contributes to the balance between maintained infection and lytic activity in tree shrew cultures. Reduced interferon-associated transcription raises a complementary question about host responsiveness. The present enrichment results nominate these relationships for functional testing but do not establish MYC/E2F1 dependence or impaired interferon signaling.

The lytic-induction experiments provide the principal functional advance of the cellular platform. Zta expression and sodium butyrate treatment are established approaches to induction (31, 32), but here they test the responsiveness of infection maintained in tree shrew cells. Zta expression increased downstream EA-D/BMRF1 and capsid p18 signals in both backgrounds, demonstrating a response beyond detection of the introduced regulator itself. Sodium butyrate increased lytic protein expression and DNase-resistant EBV DNA in culture supernatants. Preparations recovered from treated-culture supernatants generated approximately 7% GFP-positive Raji recipients for T-antigen donors and 10% for hTERT donors, providing functional evidence of infectious EBV production. Thus, maintained infection is compatible with retained lytic competence rather than being defined solely by persistent viral markers. The system therefore enables functional investigation of viral regulation after infection has been established.

Beyond experimentally induced reactivation, identifying the host signals that regulate the latent-to-lytic transition is an important next step. We recently demonstrated that IL-17A induces EBV lytic reactivation and infectious virus production in human B-cell models with latency I, identifying an inflammatory cytokine as a direct regulator of the viral life cycle (33). The dependence of this response on cellular and latency context highlights the importance of complementary experimental systems. The combination of longitudinal infection monitoring and species-matched cultures established here provides a foundation for investigating whether and how inflammatory cues influence EBV persistence and reactivation in Chinese tree shrews.

Recent independent work further supports the biological rationale for this model. A tree shrew immune-cell atlas identified human-like tonsillar structures and conserved NR1H3-positive macrophage functions, and characterized immune remodeling during acute EBV infection (34). These findings provide comparative and immunological context for the prolonged persistence and species-matched cellular systems described here. They also highlight opportunities to connect native immune-cell responses with tissue reservoirs in future studies.

The late liver and spleen EBER findings identify a focused next question: which cell populations contain EBV during persistent infection? Cell-type-resolved localization, including analysis of oral tissues, could connect tissue signals with intermittent blood and oral detection. In the cultures, the relationship between growth-associated transcription, interferon responsiveness, and viral activity provides a focused direction for subsequent work. Functional receptor studies and sample-matched host and viral profiling would further define the biology of the system. Additional independently derived cultures would help distinguish infection-associated host states from culture selection. These complementary systems support subsequent studies of the determinants of persistence and reactivation.

In conclusion, Chinese tree shrews combine evolutionary proximity to primates, an immunocompetent native host environment, and the practical scale of a small animal. This study extends the model beyond susceptibility by combining blood viral DNA detection beyond two years and late liver and spleen EBER localization with promoter-associated transcription and inducible infectious output in species-matched cultures. The resulting animal-cell platform complements humanized mice and nonhuman-primate systems for investigating persistence, viral regulation, and infection-associated host states.

## ACKNOWLEDGEMENTS

Research reported in this publication was supported by the National Institute of Allergy and Infectious Diseases as well as National Cancer Institute of the National Institutes of Health under Award Number R01AI195576 and R01CA261258 to ZL and R01CA262090 and R01CA272142 to EKF. The content is solely the responsibility of the authors and does not necessarily represent the official views of the National Institutes of Health. This work was also supported by a U.S.-Japan Cooperative Medical Sciences Program Collaborative Award from the National Institute of Allergy and Infectious Diseases and CRDF Global (grant number DAA3-19-65602-1), a Louisiana Cancer Research Center seed grant, a Carol Lavin Bernick faculty grant, and a Ladies Leukemia League research grant to ZL.

---

**Figure S1. Serial EBV DNA testing of blood and oral specimens from uninoculated Chinese tree shrews (NC01-NC06).** Blood (left panels) and oral-swab specimens (right panels) were collected from six uninoculated Chinese tree shrews (NC01-NC06) at variable intervals during the observation period and tested for EBV DNA by quantitative PCR (qPCR). Each pair of panels represents one animal, with individual specimens plotted at their respective collection times. EBV DNA was not detected in any of the tested blood or oral specimens. Open circles indicate nondetected (ND) samples and are displayed at a fixed position below the quantitative range for visualization on logarithmic axes. The ND plotting position does not represent a measured concentration or an assay detection limit. Time is expressed as days from first collection for each animal. Sampling frequency and duration varied among animals.

**Figure S2. Full-length human/tree shrew CR2 alignment.** Human CR2/CD21 NP_001006659.1 versus tree shrew XP_014447439.1: 795 identical residue pairs across 1,094 alignment columns (72.67% identity), including five gap-containing columns. Numbers use the respective full-length precursor coordinates. Shading/vertical bars indicate identity, colons positive BLOSUM62 substitutions, dots other substitutions, and dashes gaps. The targeted SCR1-SCR2 alignment is assigned to main Figure 3C.

**Figure S3. Full-length human/tree shrew EphA2 alignment.** Human EphA2 NP_004422.2 versus tree shrew TSDBTID.7937: 925 identical residue pairs across 976 alignment columns (94.77% identity), without gaps. Numbers use full-length precursor coordinates. Shading/vertical bars indicate identity, colons positive BLOSUM62 substitutions, and dots other substitutions. The targeted ligand-binding-domain alignment is assigned to main Figure 4B.

**Figure S4. Expression patterns of the 50 top-ranked exploratory host candidates.** The heatmap shows the 50 exploratory candidates with the lowest adjusted P values from the blocked DESeq2 analysis controlling for immortalization background. Ties were resolved by decreasing absolute shrunken log2 fold change. Values are variance-stabilized counts standardized within each gene across all six cultures to obtain row z scores. Rows and columns were hierarchically clustered. Column annotations indicate EBV status and hTERT or TAg background. Cntl denotes the corresponding EBV-negative parental culture. The same six RNA-seq libraries were used for Figure 7C, which displays a separately selected 24-gene high-confidence subset.

## DATA AVAILABILITY

Raw sequencing data are described in the separate Database accession number statement in Materials and Methods. Supporting results are presented in the article and supplemental material. Numerical source data and custom R/Python analysis and figure-generation scripts are available upon request. RNA-seq analyses of the tree shrew cell cultures used FastQC (https://www.bioinformatics.babraham.ac.uk/projects/fastqc/), fastp (https://github.com/OpenGene/fastp), MultiQC (https://github.com/MultiQC/MultiQC), STAR (https://github.com/alexdobin/STAR), SAMtools (https://www.htslib.org/), featureCounts (Subread) (https://subread.sourceforge.net/), Salmon (https://salmon.readthedocs.io/), BEDTools (https://bedtools.readthedocs.io/), gffread (https://ccb.jhu.edu/software/stringtie/gff.shtml). Baseline tissue virome screening used STAR and IGV (https://igv.org/). Transcript summarization, differential-expression analysis, and gene-set enrichment used R (https://www.r-project.org/), tximport (https://bioconductor.org/packages/tximport/), DESeq2 (https://bioconductor.org/packages/DESeq2/), apeglm (https://bioconductor.org/packages/apeglm/), fgsea (https://bioconductor.org/packages/fgsea/), msigdbr (https://igordot.github.io/msigdbr/). Data handling and plotting used dplyr (https://dplyr.tidyverse.org/), tidyr (https://tidyr.tidyverse.org/), tibble (https://tibble.tidyverse.org/), readr (https://readr.tidyverse.org/), ggplot2 (https://ggplot2.tidyverse.org/), ggrepel (https://ggrepel.slowkow.com/), pheatmap (https://cran.r-project.org/package=pheatmap). Custom computational analyses also used Python (https://www.python.org/), and non-transcriptomic statistical analyses used GraphPad Prism (https://www.graphpad.com/scientific-software/prism/). Gene-set, protein-annotation, and structural resources included MSigDB (https://www.gsea-msigdb.org/gsea/msigdb), UniProt (https://www.uniprot.org/), Protein Data Bank (https://www.rcsb.org/). Software versions are reported in Materials and Methods.

